# Revisiting Long-Distance Axon Growth Ability in the Developing Spinal Cord using a Novel Lesion Paradigm

**DOI:** 10.1101/2022.03.20.484375

**Authors:** Carolin Ruven, Julia Kaiser, Payal Patel, Francesca Serraino, Riki Kawaguchi, Vibhu Sahni

**Affiliations:** Burke Neurological Institute, White Plains, NY, 10605; Department of Psychiatry and Behavioral Sciences, University of California Los Angeles, Los Angeles, CA 90095; Feil Family Brain and Mind Research Institute, Weill Cornell Medicine, New York, NY 10065; Weill Cornell Graduate School of Medical Sciences, New York, NY, 10065

## Abstract

Established experimental lesions in the developing central nervous system (CNS) disrupt microenvironments critical for long-distance axon growth and guidance. Therefore, the precise developmental time at which the CNS ceases its capacity to support long-distance axon growth remains unknown. Accordingly, we established a new microsurgical approach to axotomize developing corticospinal tract (CST) axons in the neonatal spinal cord while leaving their local microenvironments relatively intact. This enables unambiguous investigation of long-distance CST growth ability in the CNS. Surprisingly, we find that CST axons lose their capacity for long-distance growth even during the developmental period of CST extension. While this ability remains intact in spinal locations where normal CST extension is occurring, it is completely abolished at sites distant from these locations. Further, the developmental time window for which this ability is maintained is much shorter than for other forms of axon growth such as sprouting. Long-distance CST growth ability does not correlate with astrocytic or microglial activation, nor with myelination levels. These results indicate that long-distance CST growth is controlled by mechanisms that operate early in development in a time- and region-specific manner.

---

One of the critical requirements of brain function is network connectivity, whereby information is transmitted among computational nodes over long distances via white matter connections ^1^. The CST exemplifies such long-range connections between the cortex and its subcortical targets, including the spinal cord. This connectivity is established during development via fasciculated axon extension in the developing white matter, which supports efficient long-distance axon navigation. Long-distance CST growth has not been achieved to date after adult injuries in the spinal cord. However, such growth does not occur even in established neonatal lesion paradigms that are traditionally thought to exhibit a high capacity for supporting axon regeneration. So, a fundamental question remains: when is this ability lost during development? A central scientific tenet regarding axon regeneration in the mammalian CNS is that regenerative ability remains high during development and declines into adulthood (reviewed in ^2^). This decline is thought to occur due to a reduction in intrinsic growth capacity of neurons ^3-6^ and developmental changes in the CNS environment ^2,7-9^. The environmental differences include differential responses of neonatal versus adult glia to injury resulting in an axon growth-restrictive environment after lesions to the adult CNS ^10,11^, although adult CNS lesions can elicit multiple, distinct forms of axonal plasticity ^12,13^.

Most long-distance axon growth and high-fidelity guidance in the CNS occurs during embryonic life, hindering experimental manipulation. The rodent CST extends into the spinal cord entirely postnatally and therefore provides the unique advantage of relatively easy experimental access to investigate long-distance axon growth competence during the normal developmental period of axon extension. Numerous investigations have identified that lesions to the developing CST in neonates (e.g., compression, over-hemisection, transection) result in greater capacity for anatomical plasticity and better functional outcomes than similar lesions in the adult ^14-22^. Seminal work in rodents identified that neonatal lesions elicit plasticity in CST trajectory, where axons are re-routed around lesions and exhibit some, although limited, growth ability into the lesion site. However, in all these instances, long-distance CST growth is largely abolished ^18,21^. A critical limitation of these lesion paradigms is that they cause significant disruption to the spinal cord neuronal, glial, and vascular microenvironments, which play a critical role in normal developmental axon growth and guidance ^23^. This therefore precludes the use of such lesions to investigate the capacity of the CNS to foster long-distance axon growth and guidance. For instance, an over-hemisection at thoracic T8-10, even prior to the arrival of CST axons, resulted in only a few axons extending into the lesion and no long-distance growth ^21^, indicating that the normal developmental process of axon extension is largely disrupted.

To overcome this issue and determine when the ability for long-distance CST axon growth is lost, we established a novel approach to axotomize the developing CST without causing overt spinal damage. This approach enabled precise delineation of the spatiotemporal trajectory of the loss of long-distance CST growth ability through development in the absence of measurable changes in the cellular environment of the spinal cord. We find that long-distance CST growth ability closely parallels the normal developmental trajectory of CST extension into the cord in the spinal white matter. Our results indicate that this ability is lost on the time scale of days. We identify that when CST axons first arrive at a given spinal segment, there is a brief window of ∼4-5 days when that spinal level retains the capacity to support long-distance CST growth, after which this ability is lost. Therefore, long-distance CST growth ability is lost at distinct times at distinct spinal levels during development. Further, the ability for other forms of axonal growth such as sprouting is lost at distinct developmental times. Even when CST axons lose their ability for fasciculated, long-distance growth in the white matter, they still retain the ability for growth in the gray matter; a form that appears to resemble regenerative sprouting. Finally, the loss of long-distance CST growth at distinct spinal segments does not correlate with differences in levels of astrocytic or microglial reactivity nor with differences in levels of myelination. Our results suggest that context-specific axon guidance mechanisms that operate on short time scales represent an earlier control over long-distance axon regenerative ability before more global regulators, such as the intrinsic growth capacity of neurons and the changes in the environment, limit all forms of axonal plasticity during development. This experimental approach now provides a novel paradigm to investigate potential mechanisms controlling long-distance axon regeneration in the mammalian CNS.

## Newly established microsurgical lesions enable CST axotomy with minimal damage to the spinal environment

Established experimental models of neonatal spinal cord injuries in rodents cause significant damage to the spinal cord, disrupting the guidance cues that normally direct CST axons in development. This limits the ability of such models to interrogate the competence of the CNS to support long-distance CST growth. We therefore established a new microsurgical approach to axotomize the developing CST with a beveled glass micropipette vibrating at an ultrasonic frequency under visual guidance provided by ultrasound-guided backscatter microscopy (Extended Data Fig. 1a-e, Extended Data Video 1). These microsurgical lesions (hereby referred to as “microlesions”) axotomize the dorsal funiculus and cause minimal overt damage to the surrounding tissue leaving the spinal environment largely unperturbed (whole mount view of the spinal cord at P35 after a P4 microlesion is shown in Extended Data Fig. 1f). Immunohistochemistry for glial fibrillary acidic protein (GFAP) and ionized calcium binding adaptor molecule 1 (Iba1) shows minimal astrocytic and microglial reactivity, respectively at the microlesion (Extended Data Fig. 1g -g”) (we later perform more in-depth analyses of astrocytic and microglial reactivity in Fig. 3, Extended Data Fig. 5, as well as transcriptomic analyses indicating that the broad topography of the spinal cord is maintained after microlesions; Extended Data Fig. 10). Together, these data indicate that microlesions cause minimal overall damage to the spinal cord.

### Long-distance CST growth is lost at distinct times at distinct spinal levels

Using our novel microlesions, we first asked whether CST axons, lesioned during the developmental period of axon extension into the cord, are still able to maintain competence for long-distance growth. In mice, the CST extends into the spinal cord during the first postnatal week. At postnatal day 4 (P4), pioneer CST axons are traversing the caudal-most thoracic segments ^24-26^, which suggested that long-distance CST growth ability would be maintained in the caudal thoracic cord. We therefore first investigated long-distance CST growth ability following P4 microlesions at thoracic T11. To visualize CST axons, we delivered AAV-tdTomato into cortex (Fig. 1f) and analyzed CST extension at P35. We quantified CST extension past the microlesion in both, the dorsal funiculus (DF) (normal location for majority of CST axons in rodents), and the dorsolateral funiculus (dLF) (where only a minority of CST axons normally transverse) (Fig. 1e, g, Extended Data Fig. 2). Following a P4 microlesion at T11, we find that long-distance CST growth in the DF is indistinguishable from non-lesioned controls. In non-lesioned control mice, 10 ± 3% of CST axons present in the ventral medulla reach lumbar L2. Similarly, following a P4 microlesion at T11, 9 ± 2% of CST axons reach L2 (Fig. 1e, h-i, right panels). We confirmed these findings using optically cleared spinal cords (Extended Data Fig. 3, Extended Data Video 2). This indicates that the CST maintains robust long-distance growth competence at thoracic T11 during the period of normal CST growth at this spinal level. Further, our results also indicate that microlesions do not significantly disrupt the extrinsic spinal environment and that the minimal astrocytic and microglial reactivity induced by microlesions (further shown in Fig. 3 and Extended Data Fig. 5) does not interfere with the normal developmental processes of CST axon growth and guidance.

**Fig. 1.**
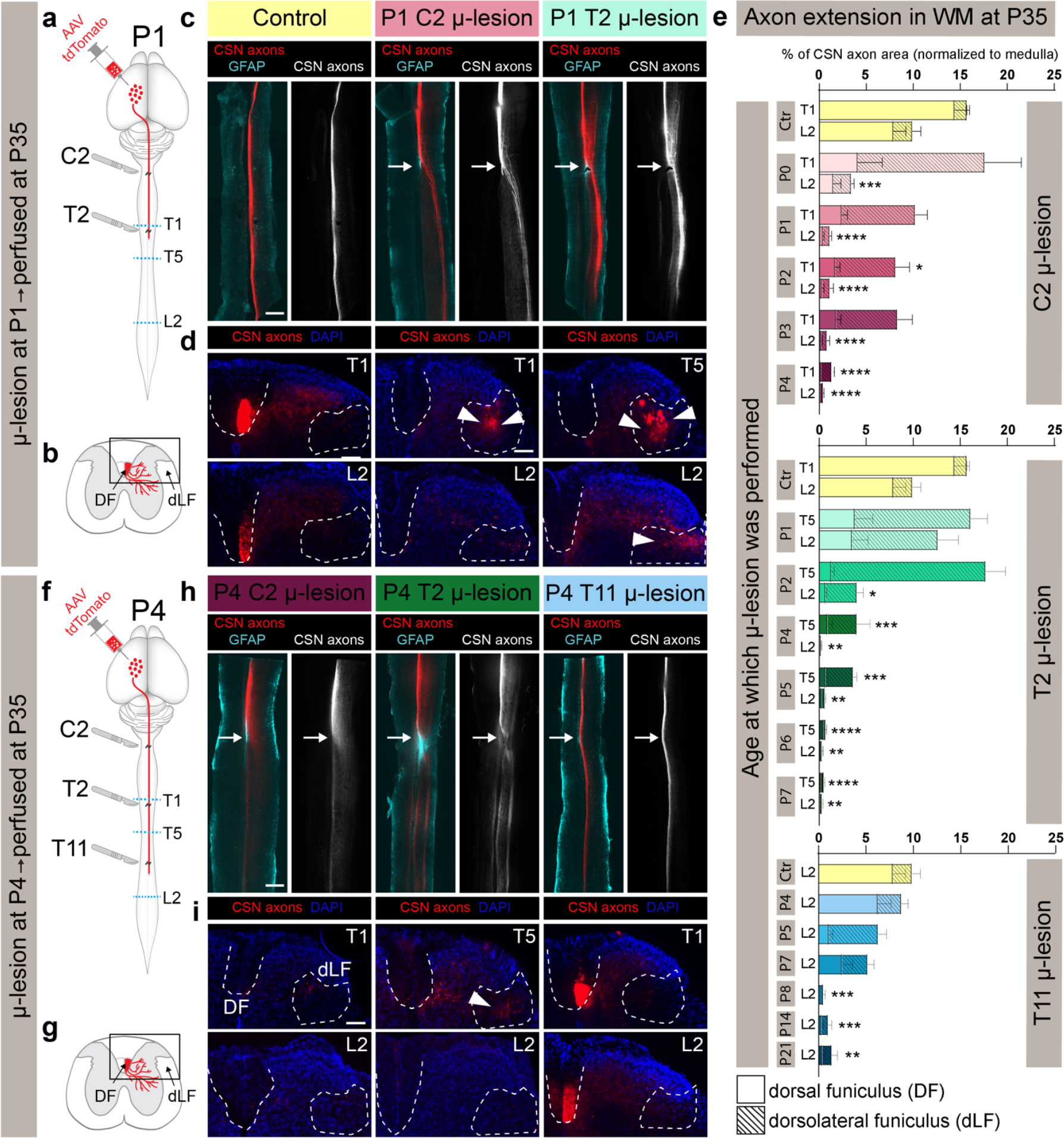
Segmentally distinct CSN axon growth responses to microlesions at distinct spinal levels. **a, b, f, g**. Schematics outlining experimental protocol and analysis. **a, f**. AAV-tdTomato was delivered into caudomedial cortex and mice underwent either a P1 or P4 microlesion at one of three distinct spinal segmental levels (C2, T2, or T11 as indicated by scalpels). The segmental level where growing CST axons are present at the time of the microlesion is indicated by the level of the red line in schematics. Mice were analyzed at P35. Blue dotted lines indicate the levels at which axial sections of the spinal cords were analyzed to investigate CSN axon extension. **b, g**. Schematics of spinal axial sections showing locations in the dorsal funiculus (DF) and dorsolateral funiculus (dLF) where axon extension was quantified. **c, h**. Horizontal spinal cord sections at P35 after P1 (**c**) and P4 (**h**) microlesions at distinct spinal levels. GFAP+ astrocytes (cyan) delineate microlesion (white arrow). CST axons are in red in merged images and in monochrome. **d, i**. Axial sections of the same spinal cords caudal to the microlesion showing CSN axons either in DF or dLF (demarcated by dotted outlines). Arrowheads indicate the diverted CSN axons in the dLF after P1 microlesions at C1 and T2, as well as after P4 microlesions at C2. **e**. Quantification of CSN axon area at thoracic T1/T5 and lumbar L2 at P35 following a microlesion at either cervical C2, thoracic T2 or thoracic T11 at distinct postnatal ages. The total (DF + dLF) CSN axon area is plotted; hatched bars indicate axon area quantified in dLF. For individual quantification of DF and dLF, see Extended Data Fig. 2. ** p<0.01, *** p<0.001, **** p<0.0001 control DF+dLF vs. microlesioned DF+dLF. All data shown are mean ± s.e.m.. Scalebars: **c, h:** 500μm; **d, i:** 100μm

We next performed P4 microlesions at 2 distinct spinal segments (cervical C2, and thoracic T2) and similarly analyzed long-distance CST growth at P35 (Fig. 1f). These microlesion sites are closer to CSN soma than the microlesion at T11, and further removed from the growing ends of the CST– the C2 microlesion is farther away from the growing ends than the T2 microlesion. Given that at least some CSN are in a state of active axon growth to caudal thoracic and lumbar segments, we expected to find that long-distance CST growth competence at these more rostral spinal segments would be similarly maintained as we observed at thoracic T11. Surprisingly, we find that this is not the case. After a P4 microlesion at cervical C2, there is almost a complete loss of long-distance CST growth. In non-lesioned mice, 16 ± 2% of CST axons present at the ventral medulla extend to thoracic T1 with 10 ± 3% extending to lumbar L2. In striking contrast, following a P4 microlesion at C2, only 1.3 ± 0.3% of CST axons in the ventral medulla reach T1 (DF + dLF combined; with only 0.2% at DF), and 0.4 ± 0.1% reach L2 (Fig. 1e, h-i, left panels). Hence, >90% of CST axons failed to extend to the thoracic levels and >95% of axons failed to reach the lumbar cord. We additionally confirmed these findings in an optically cleared spinal cord, which identified that most tdTomato+ CST axons do not extend past the microlesion (Extended Data Fig. 3, Extended Data Video 2). This indicates a significant decline in long-distance CST growth ability in the cervical cord even during the period of active CST extension into thoracic and lumbar segments. When the P4 microlesion was performed at T2, i.e. closer to the growing CST ends (the CST is growing toward caudal thoracic segments at P4 ^24-26^), we observed limited long-distance CST growth (4.0 ± 1.5% of axons at ventral medulla extend to T5, but almost none extend to L2, Fig. 1e, h-i); however, these CST axons are diverted from DF to dLF (Fig. 1e, h-i, middle panel; Extended Data Fig. 2). This suggests that some CST axons retain the ability for long-distance growth even when they are not extending in their normal location in the DF. These results indicate that the ability of the CST for long-distance growth is not lost uniformly across the spinal cord. Further, this loss of long-distance CST growth ability at a given spinal level appears to depend on the proximity of the microlesion to the growing CST ends. When CST axons are axotomized closer to the growing ends there is more robust long-distance growth, and this ability declines with increasing distance between the growing ends of the CST and the site of the axotomy. This suggests that the relative distance of the axotomy from the growing ends of the CST predicts whether long-distance CST growth ability would remain intact.

To further test this idea, we next performed microlesions at P1, at a time when growing CST axons have reached thoracic T3 ^24^. We performed P1 microlesions at two distinct spinal segments (C2 and T2) and similarly analyzed axon extension at P35 (Fig. 1a). Our results are that after P1 microlesions at thoracic T2 i.e., close to the growing ends of the CST at T3, there is robust long-distance CST axon growth (16.1 ± 2.4% of axons reaching T5, and 12.6 ± 2.7 % reaching L2), which is statistically indistinguishable from non-lesioned controls. After P1 microlesions at cervical C2, CST axons exhibit long-distance growth, which is distinct from P4 microlesions at this level; however, CST axons after a P1C2 microlesion do not fully extend to their normal targets – 10.2 ± 1.6% of axons at the ventral medulla reach T1, which is slightly reduced from 15.8 ± 1.7% in controls, but only 1.1 ± 0.3% reach L2 (this is still higher than 0.4% observed after a P4 microlesion). These results indicate that overall, there is greater ability for long-distance CST growth at cervical C2 at P1 than at P4 (Fig. 1c-e). Therefore, similar to P4 microlesions, when the axotomy is performed close to the growing ends of the CST, there is more robust long-distance axon growth. However, in both these P1 microlesion groups, the majority of CST axons are diverted from DF to dLF (Fig. 1c-e, Extended Data Fig. 2). This further indicates that long-distance CST growth does not require axons to traverse in their normal location in the DF. CST axons that get diverted to the dLF, where only a minority of CST axons normally traverse, are still able to maintain some ability for long-distance growth. However, even this “diverted” long-distance growth at C2 is absent by P4, i.e., long-distance CST growth ability in the white matter at C2 is lost by P4. We next sought to establish the precise time course of the decline in long-distance CST growth ability at these distinct spinal levels – C2, T2, and T11.

We therefore performed microlesions at multiple additional time points: P0, P2, and P3 at cervical C2; P2, P5, P6, and P7 at thoracic T2; P5, P7, P8, P14, and P21 at thoracic T11. We find that long-distance axon growth is almost intact after P0 C2 microlesions, similar to P1 T2 and P4 T11 microlesions (Fig. 1e), i.e., when the microlesion occurs around the spinal segmental level that harbors leading CST ends. At cervical C2, reduced long-distance CST growth ability remains at both P2 and P3, and the ability is lost by P4. At thoracic T2, reduced ability remains at P2, and P5, while at T11 reduced ability is present at both P5, and P7 (Fig. 1e). One day later, by P6 at thoracic T2 and by P8 at thoracic T11, long-distance CST growth ability is abolished at these spinal levels, similar to the effects of P4 microlesions at cervical C2 where we find almost no axons extend across the microlesion (Fig. 1e; for mouse numbers in all the distinct groups, see Extended Data Table 1). Together, our findings indicate that long-distance CST growth ability is not lost uniformly across the spinal cord. Rather this ability is lost at distinct times at distinct levels–P4 at C2, P6 at T2, and P8 at T11. These results collectively indicate that long-distance CST growth ability closely follows the normal developmental trajectory of CST extension in the spinal cord – once the leading ends of the CST arrive at a specific spinal level, there is a window of ∼4-5 days when long-distance growth ability in the white matter is maintained; after which period this ability declines sharply (summarized in Extended Data Fig. 4). Further, for this initial decline in long-distance CST growth ability the proximity of the axotomy to the growing ends of the CST takes precedence over the proximity to the cell body. For instance, at C2, while the distance of the axotomy from the CSN soma hasn’t changed from P0 to P4, yet there is a striking difference in long-distance growth ability at these two times – the ability is intact at P0 and is lost at P4. What differs at these two times, is that the location of the growing CST – the axotomy is closer to these ends at P0 and significantly farther at P4.

### Segmentally distinct loss of long-distance axon extension even by CSN that are in a state of active axon growth

The complete lack of long-distance CST growth following a P4 microlesion at cervical C2 is surprising, since thoraco-lumbar projecting CSN (CSN_TL_) axons are still extending toward thoracic and lumbar spinal segments at this time. CSN_TL_ reside in medial sensorimotor cortex, interdigitated with bulbar-cervical-projecting CSN (CSN_BC-med_) ^25^. Therefore, with AAV-mediated anterograde labeling, axons from both cervical- and thoraco-lumbar projecting CSN are labeled in the spinal cord. At P4, while CSN_TL_ axons are extending to caudal levels of the spinal cord, cervical-projecting CSN are already collateralizing into the cervical spinal gray matter ^24^. Since the initiation of synapse formation is known to cause a decline in regenerative ability ^27,28^, this could account for some reduction in long-distance CST growth by P4 at cervical C2. It therefore remained theoretically possible that if only CSN_TL_ axons were selectively analyzed, which are still actively extending in the thoraco-lumbar spinal cord at P4, these axons might exhibit a greater ability for long-distance growth after a P4 microlesion at C2. To address this, we utilized intersectional genetic reporter mice: Crim1CreERT2;Emx1FLPo;Ai65 (CERai65 mice, ^25^). *Crim1* expression prospectively identifies CSN_TL_ during development ^25^. In these intersectional genetic reporter mice, CSN_TL_ axons can be visualized in the spinal cord via tdTomato expression (Fig. 2a) ^25^. We used these mice to specifically investigate whether CSN_TL_ axons exhibit greater long-distance growth ability when compared to the overall population of CSN axons arising from medial sensorimotor cortex. We performed P4 microlesions in CERai65 mice at either cervical C2 or thoracic T11 and analyzed long-distance axon growth as described above. While long-distance CSN_TL_ axon growth is completely intact following a P4 microlesion at thoracic T11 (14.3 ± 1.5%, versus 15.7 ± 4.3% in Control, DF + dLF in L2; statistically indistinguishable from controls); however, long-distance CSN_TL_ axon growth is almost absent following a P4 microlesion at cervical C2 (2.7 ± 0.9%, versus 26.5 ± 5.1% in Control, DF + dLF in T1) (Fig. 2b-c). These results are nearly identical to our previous findings using AAV-mediated anterograde labeling. Thus, CSN_TL_ axons lose their long-distance growth ability at cervical C2 even while they are extending toward caudal thoracic and lumbar spinal segments; this growth ability remains fully intact at thoracic T11 at the same developmental time.

**Fig. 2.**
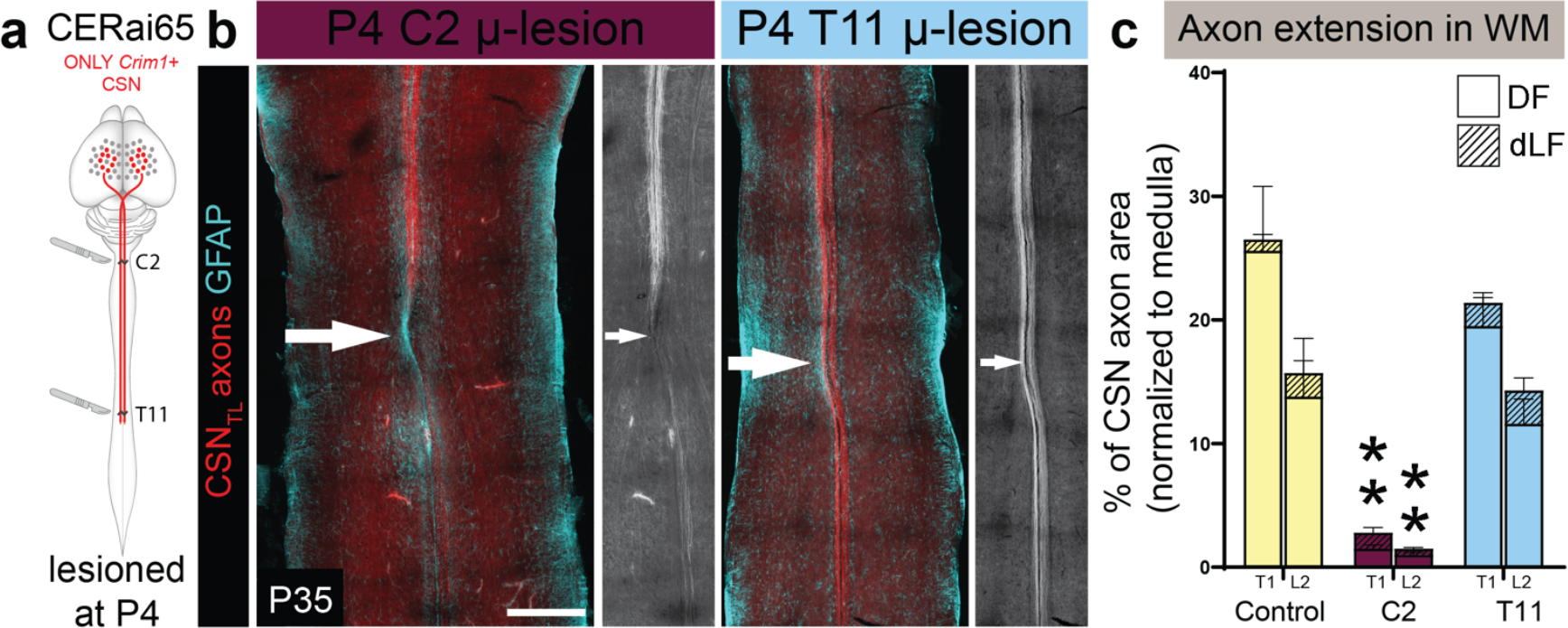
Segmentally distinct loss of long-distance axon growth occurs even by CSN that are in a state of active axon growth. **a**. In Crim1CreERT2;Emx1FlpO;Ai65 triple transgenic (CERai65) mice, only *Crim1*+ cortical neurons are labeled via tdTomato expression. *Crim1* expression identifies thoraco-lumbar projecting CSN (CSN_TL_) ^25^. P4 microlesions were performed at either C2 or T11 (indicated by scalpels in schematics). **b**. Horizontal spinal sections from CERai65 mice after P4 microlesions at either cervical C2 or thoracic T11. CSN_TL_ axons are in red and in monochrome. White arrow indicates the microlesion site. Minimal astrocytic activation is seen at the microlesion (GFAP, cyan). **c**. Quantification of CSN_TL_ axonal area extending to thoracic T1 and lumbar L2, normalized to axonal area in the ventral medulla. DF – dorsal funiculus, dLF – dorsolateral funiculus. Data are mean ± s.e.m.. ** p<0.01, control DF+dLF vs. microlesioned DF+dLF. n= 4 for Control and T11 group, n=6 for C2 group. Scalebar: **b:** 500 μm.

### Segmentally distinct loss of long-distance axon growth is not due to segmental differences in astrocytic or microglial activation

We next investigated whether the segmentally distinct effects on long-distance axon growth could be accounted for by differences in levels of astrocytic or microglial reactivity in response to microlesions. Although microlesions do not result in the formation of a classical lesion core, there is still some, albeit minimal, astrocyte reactivity. To test whether segmental differences in levels of astrocytic activation correlate with spinal level effects on long-distance CST growth, we used GFAP immunohistochemistry in P35 spinal cords to evaluate the astrocytic response to microlesions performed at distinct spinal levels at distinct developmental times (the same mice where we quantified long-distance CST growth in Fig 1e; summarized in Extended Data Fig. 4). For these analyses we used serial horizontal sections to reconstruct the extent of the microlesion across the entire volume of the spinal cord. Representative images from mice across all P1 and P4 microlesioned groups show no overt difference between any of the microlesions (Fig. 3a). We quantified GFAP intensity at the microlesion and find no difference between microlesion groups that either foster or do not support long-distance axon growth (Fig. 3b). We also performed 3D volumetric reconstructions of the subtly increased GFAP immunoreactivity at the microlesion. Representative 3D reconstructions from mice that underwent P4 microlesions at C2, T2, and T11 show that this volume is equivalent across all 3 mice, even though CSN axons show very distinct long-distance growth responses across these microlesion sites– there is robust long-distance growth through the P4 T11 microlesion, reduced, but diverted growth through the P4 T2 microlesion, and no growth through the P4 C2 microlesion (Fig. 3c). Quantification of the 3D volume of increased GFAP immunoreactivity also finds no difference between all microlesioned groups (Fig. 3d). We also measured the area of increased GFAP+ reactivity in serial sections of the spinal cord at P35 across 5 different microlesioned groups– P1 C2, P1 T2, P4 C2, P4 T2, and P4 T11. These results also find that the spatial distribution of increased GFAP immunoreactivity across distinct microlesions does not differ; the area of increased reactivity extends ∼800 μm rostrocaudally and ∼300 μm mediolaterally across all groups (Extended Data Fig. 5). We similarly quantified microglial responses using Iba1 immunohistochemistry and find that the spatial distribution of microglial reactivity is strikingly less than that observed for astrocytes; further, and more importantly, there is no difference between the groups at P35 (Extended Data Fig. 5). These results highlight a very minimal glial response to the microlesion.

**Figure 3.**
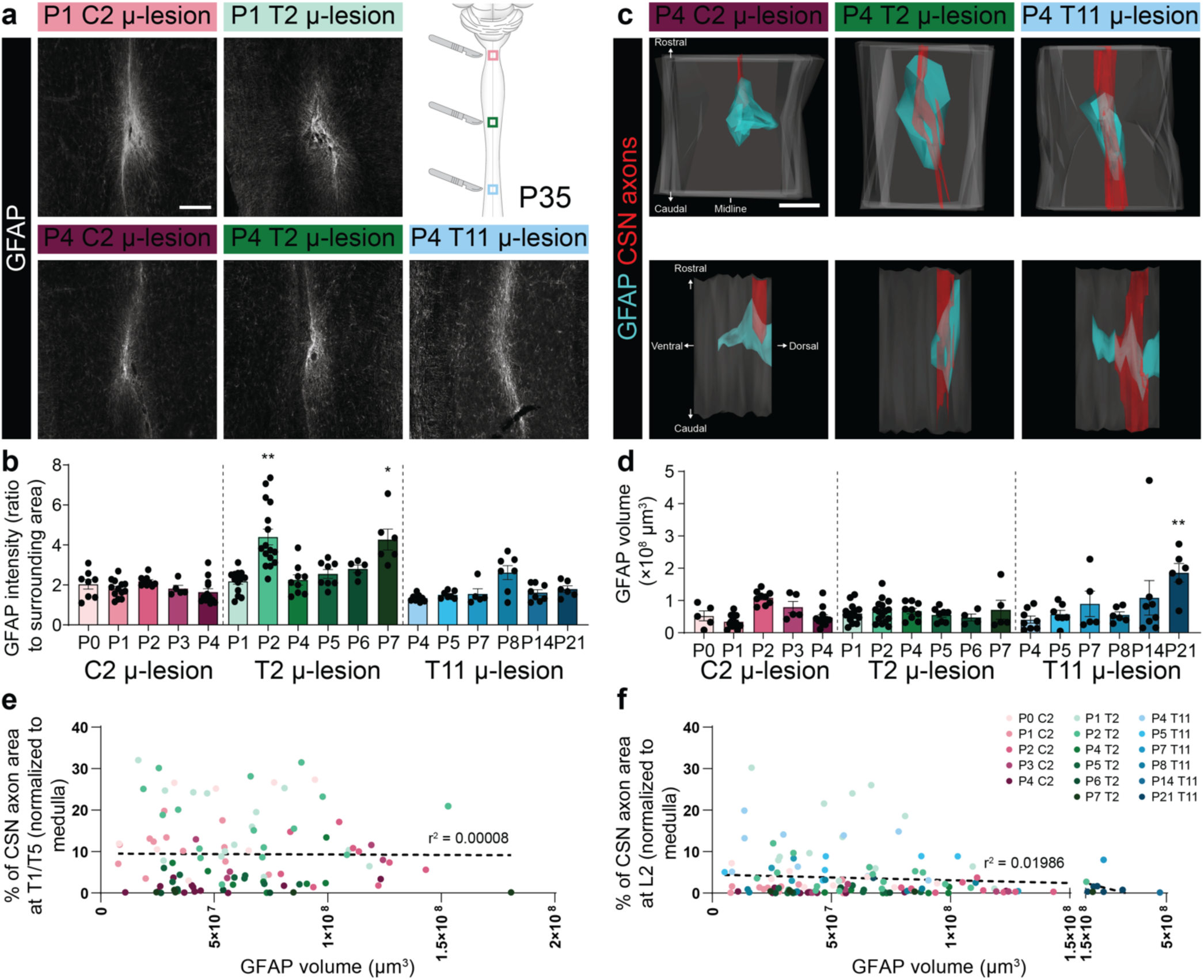
Segmentally distinct loss of long-distance CSN axon extension is not due to segmental differences in astrocytic activation. **a**. Representative images of GFAP immunohistochemistry (astrocytic response) on horizontal sections of the mouse spinal cord at P35. Magnified views of the microlesion site are shown following either P1 or P4 microlesions at the distinct spinal levels indicated. The schematic on the top right shows the spinal locations for the images. **b**. Quantification of GFAP intensity at the microlesion site across different microlesioned groups. There is no significant difference between the multiple, distinct groups. **c**. 3D reconstructions of the microlesion site (GFAP in cyan, CSN axons in red) from the ventral view (upper row) and lateral view (lower row). **d**. Quantification of volume of increased GFAP reactivity across the different microlesioned groups. ** p<0.01, P21 T11 microlesion group is significantly different from the following groups: P0, P1 and P4 C2 groups; all T2 groups; P4, P5, and P8 T11 groups. **e-f**. Pearson correlation between GFAP volume at microlesions and CSN axon growth to thoracic (**e**) or lumbar (**f**) cord. Each dot represents a single mouse, color-coded by its microlesion group (color code on top right). GFAP volume does not correlate with axon growth. All data shown are mean ± s.e.m.. Scalebars **a**: 200 μm; **c**: 500 μm.

Finally, we performed correlation analyses between the volume of GFAP reactivity at a microlesion site and the ability of that microlesion site to support long-distance axon growth to either thoracic (Fig. 3e) or lumbar levels (Fig. 3f). We find no correlation between the volume of increased GFAP+ immunoreactivity at the microlesion sites and ability for long-distance CSN axon growth after microlesions at those segmental levels.

Together, these results indicate that the differential loss of long-distance CST growth ability across developmental space and time does not arise from differences in astrocytic or microglial activation.

### Different forms of axonal plasticity are lost at distinct developmental times

In addition to long-distance growth, axons can exhibit multiple forms of plasticity, such as sprouting ^13^. We therefore investigated whether the ability for CST sprouting is lost at the same time as the ability for long-distance CST growth during development. Our results identified that at P4 the white matter at cervical C2 had lost the ability to foster long-distance CST growth. We therefore investigated whether CST axons retained the ability for other forms of growth at this time by analyzing the response of lesioned CST axons after P4 microlesions at cervical C2. At P7, three days following the microlesion, we find no outgrowth from the axotomized CST. However, by P15, we find labeled, tdTomato+ CST axon collaterals in the cervical gray matter. This collateral sprouting extends unilaterally well past the level of the microlesion with a few collaterals reaching caudal cervical spinal segments (Extended Data Fig. 6). By P35, the density of these collaterals increases significantly such that there is a high density even at caudal cervical segments (Extended Data Fig. 6); however, these collaterals do not extend into the thoracic cord (as shown in axial sections in Fig. 1e). These results indicate that CST axons retain some ability for sprouting in the gray matter, at a time when the ability for long-distance growth in white matter is abolished. We tested this idea further by investigating when CST axons lose their ability for gray matter sprouting at a given spinal level. For this, we performed T11 microlesions at P8, when long-distance axon growth is abolished by P8 (Extended Data Fig. 7; quantified in Fig. 1e). Following these P8 microlesions, while long-distance CST growth in the spinal white matter was abolished, tdTomato+ CST collaterals still extend past the lesion into the thoracic gray matter caudal to the microlesion (assessed at P35, Extended Data Fig. 7a). However, when T11 microlesions were performed at P14, there is a striking reduction in the number of gray matter collaterals caudal to the microlesion indicating a loss of both, long-distance axon growth and collateral sprouting in the gray matter (Extended Data Fig. 7b). Thus, the ability for long-distance axon growth in the white matter is lost prior to the ability for axonal sprouting.

### Mechanisms controlling segmentally distinct loss of long-distance CST growth are in effect acutely after microlesions

The results detailed thus far were obtained at P35, i.e., more than one month after the microlesions were performed. Therefore, there remained the possibility that in instances where we observed no long-distance growth at P35, CSN axons might have initiated long-distance growth in the few days after the microlesion that was subsequently pruned (before P35). This would give the appearance of no long-distance axon growth when the spinal cord was analyzed at P35. We therefore investigated CST axons acutely after P4 microlesions at distinct spinal levels– C2, T2, and T11– by analyzing the spinal cords 72 hours later. We find that the distinct effects on long-distance CST growth at these three spinal levels observed at P35, are evident 3 days after microlesions. While axons have already fully extended past the microlesion at T11, CST axons have not extended past the microlesion at C2. In addition, CST axons have already diverted into the dLF after the microlesion at T2. Notably, CST axons can be seen extending past a site of GFAP activation at the microlesion at thoracic T11 (Fig. 4a). We find no overt differences in GFAP immunoreactivity between microlesions at these distinct spinal levels (Fig. 4a), which again indicates that these distinct effects on long-distance CST growth are not due to differences in astrocytic activation. We quantified axon extension in DF and dLF, and find that the observed differential effects on long-distance axon growth occur within the first 72 h after microlesions (Fig. 4b). Further, the absence of long-distance CST growth following P4 microlesions at cervical C2 observed at P35 is not due to an initial regenerative attempt that is then aborted. This also suggests that mechanisms controlling long-distance CST growth ability are in effect acutely after the microlesion.

**Figure 4.**
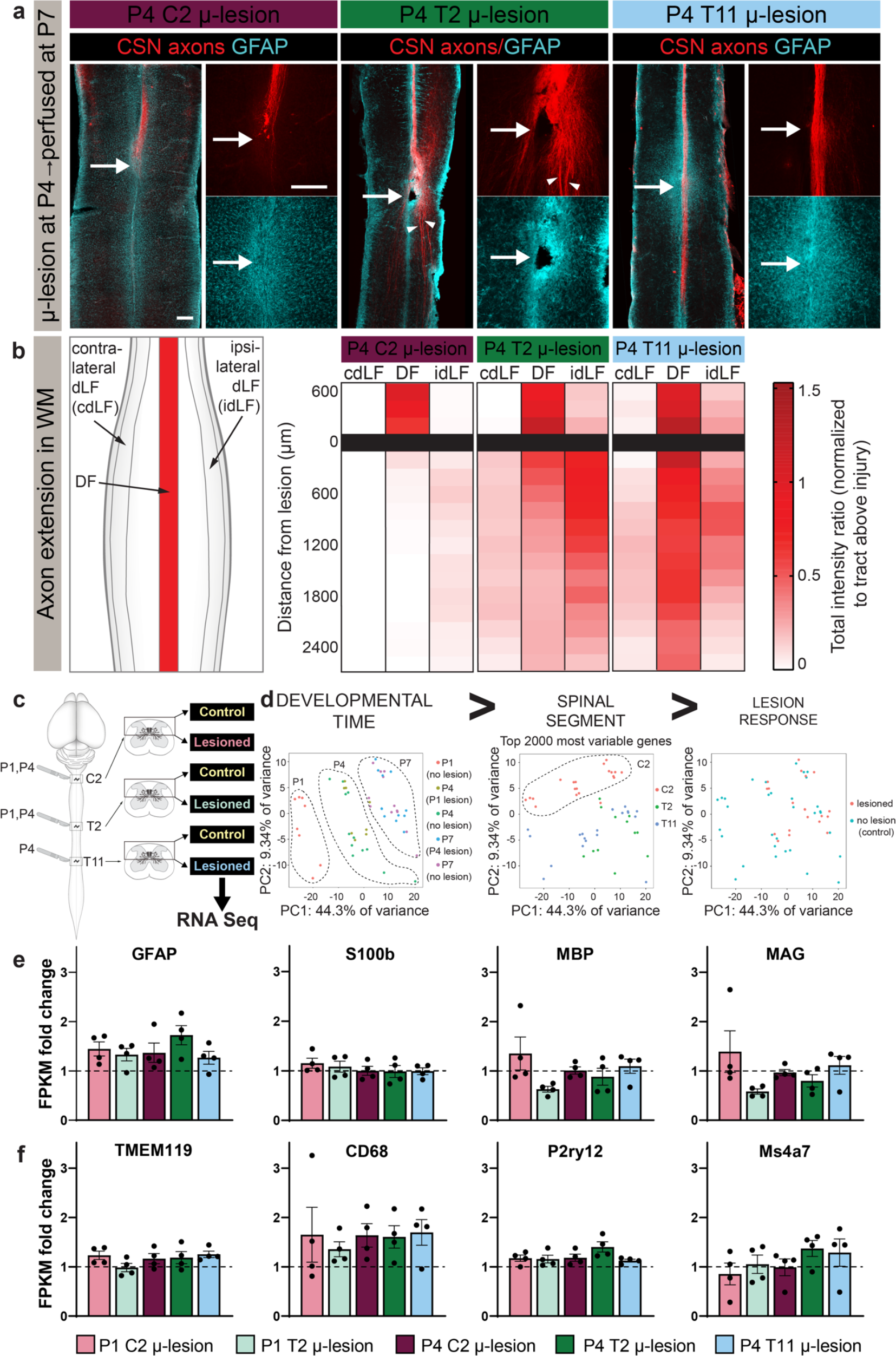
Transcriptomic analyses find similar levels of overall astrocytic and microglial activation as well as myelination at distinct spinal levels with little effect from microlesions. **a**. Horizontal sections of P7 mouse spinal cords after P4 microlesions at either C2, T2, or T11, analyzed 72 hours after microlesion. CSN axons are in red and GFAP+ astrocytes in cyan. Note that the segmentally distinct axon growth responses at distinct spinal levels are already evident at this acute time after microlesions. CSN axons do not extend past the C2 microlesion but do extend past the T11 microlesion. CSN axons can be seen diverted to the dLF after a T2 microlesion (arrowheads). **b**. Quantification of CST extension individually in the dorsal funiculus (DF), as well as in the ipsilateral/contralateral dorsolateral funiculus (idLF, cdLF, respectively). n=5 for each group. **c**. Schematic of experimental outline: RNA was extracted from the dorsal cord at C2, T2, or T11 from non-lesioned controls at P1, P4, and P7. In addition, RNA was extracted 3 days after the following microlesions – P1 C2, P1 T2, P4 C2, P4 T2, and P4 T11. **d**. Multi-dimensional scaling (MDS) plots showing that developmental time, indicated by principal component 1 (PC1) on the x-axis, is the top contributor to the variance accounting for 44.3% of the total variance across groups. PC2 on the y-axis distinguishes segmental differences accounting for 9.3% of the variance. Microlesions produce minimal gene expression changes with no contribution toward the top 2000 differentially expressed genes between the groups. **e**. Gene expression for markers of astrocytes (GFAP, S100b) and myelin (MBP, MAG), presented as fold change over non-lesioned controls (calculated from FPKM). **f**. Expression of microglial genes that promote scar-free healing in the neonatal CNS (Li, et. al., 2020 *Nature* ^10^); Tmem119 – microglia, Cd68 – activated macrophages or microglia, P2ry12 – homeostatic microglia, Ms4a7– embryonic microglia. Dotted horizontal line indicates a fold-change of 1. There are no differences in expression levels between any of the microlesioned groups. All expression data are fold change of FPKM and presented as mean ± s.e.m.. n=4 for each group. Scalebar in **a:** 500 μm.

### Transcriptional profiling indicates that segmentally distinct effects on long-distance CSN axon growth are not due to acute differences in astrocytic or microglial activation, or myelination levels

Given that the segmentally distinct effects on long-distance CST growth at P4 occur within 72 hours after the microlesion, we first investigated whether this segmentally distinct loss of long-distance CST growth at distinct spinal levels is due to segmental differences in the numbers of astrocytes, oligodendrocytes, or microglia in the spinal white matter (i.e., in the dorsal funiculus) at the time when these microlesions were performed. We therefore analyzed P4 spinal cords in non-lesioned mice at cervical C2, thoracic T2, and thoracic T11, using immunohistochemistry— GFAP (astrocytes), MBP (oligodendrocytes), and Iba1 (microglia). We find no overt differences between these distinct spinal segments (Extended Data Fig. 8).

To identify potential regulators over the early and segmentally distinct loss of long-distance CST growth ability, we performed transcriptional profiling of the dorsal spinal cord with or without microlesions at distinct spinal levels at defined developmental times (schematized in Fig. 4c). We extracted RNA from non-lesioned controls at C2, T2, and T11 at 3 developmental times – P1, P4, and P7. Also, since the segmental differences in long-distance CST growth are apparent within 3 days of the different microlesions, we collected RNA 3 days after the following microlesions: P1 at C2, P1 at T2, P4 at C2, P4 at T2 and P4 at T11. We first analyzed the non-lesioned groups that spanned multiple spinal segments at distinct developmental times (Extended Data Fig. 9). As expected, we find increasing levels of astrocytic (GFAP, S100b) and myelin (MBP, MAG) genes from P1 to P4 to P7 across all spinal levels. Interestingly, we didn’t find differences in expression levels of any microglial genes across these time points. Consistent with our immunohistochemical analyses (Extended Data Fig. 8), at any given developmental time, there wasn’t a significant difference in the levels of any of these macro- and microglial genes, across the different spinal segments. Importantly, the expression levels of these genes at a given spinal segment does not predict whether that segment can foster long-distance CST growth. At any given developmental time, spinal segments that either support long-distance growth or not (indicated by √ or X in the bar graph in Extended Data Fig. 9) can have identical expression levels of these genes (e.g., P4 C2 and P4 T11). These transcriptomic analyses extend and confirm our histological findings that there is no correlation between long-distance CST growth following a microlesion at a given spinal level and the astrocytic reactivity at that microlesion site (as shown in Fig. 3e, f).

To begin to elucidate mechanisms that might underlie this early loss of long-distance CST growth, we next performed differential gene expression analysis across all the groups and then used multi-dimension scaling to compare the differences (8 non-lesioned samples: P1 C2, P1 T2, P4 C2, P4 T2, P4 T11, P7 C2, P7 T2, P7 T11; 5 microlesioned samples with microlesions at P1C2, P4C2, P1T2, P4T2, and P4T11). Using the top 2000 differentially expressed genes, we identified distinct principal components (PC) that contribute to these molecular differences – PC1 accounts for nearly 45% of the variance and distinguishes molecular differences across developmental time, while PC2 accounts for ∼10% of the variance and distinguishes segmental differences (multidimensional (MDS) plot showing these PCs in Fig. 4d). Hence developmental time (P1 vs. P4 vs. P7) is the greatest contributor to molecular differences between groups, followed by the spinal segmental level (Fig. 4d).

We next investigated the effect of the microlesions on the molecular architecture of the spinal cord. Consistent with the small extent of the microlesions, we find minimal differences between non-lesioned controls and the corresponding microlesioned samples (Fig. 4d). Therefore, molecular differences between distinct spinal segments, even in non-lesioned mice, are far greater than the effects of the microlesions themselves. Consistent with this, we also analyzed the 5 different microlesioned groups with their corresponding non-lesioned controls to see if the broad topography of the spinal architecture is maintained after microlesions. We find no significant differences in expression levels of broad neuronal or cytoskeleton genes, dorsal cord transcription factors, nor in levels of AMPA and NMDA receptor genes (Extended Data Fig. 10), indicating that microlesions cause minimal disruption to the spinal cord. Next, we investigated whether the microlesions at distinct spinal levels elicited differential responses from the principal glial cell types that might account for the distinct effects on long-distance axon growth. We therefore analyzed the 5 different microlesioned groups for fold-changes in levels of astrocytes, microglia, and myelin genes as compared to their non-lesioned counterparts. Once again, consistent with our histological analyses, RNA-sequencing finds no differences in GFAP (astrocytes), S100b (astrocytes), MBP (myelin basic protein), and Mag (myelin-associated glycoprotein) across all 5 groups (Fig. 4e). We also analyzed microglial genes that are now known to promote scar-free healing in the neonatal spinal cord ^10^ to investigate whether these genes are differentially activated following microlesions across all 5 microlesioned groups. We similarly find no difference in the fold-change in any of these genes across the different groups (Fig. 4f). Together these results indicate there are no broad differences in the responses of these principal glial cell types to microlesions across distinct spinal levels at different developmental times. These data further indicate that the segmentally distinct effects on long-distance CST growth ability do not correlate with differences in astrocytic or microglial activation or segmental differences in levels of myelination.

Developmental time and segmental levels were the principal contributors to the variance across the groups (Fig. 4d). Therefore, to identify potential regulators that might support or inhibit long-distance CST growth regardless of developmental time or segmental level, we next compared only the microlesioned groups, based on whether that spinal level supported robust, moderate, or no long-distance CST growth as identified by our anatomical findings. We therefore classified the 5 microlesioned samples into the following three groups: 1) no long-distance CST growth (P4 C2); 2) moderate long-distance CST growth (P1 C2, P4 T2); and 3) high long-distance CST growth (P1 T2, P4 T11) (Extended Data Fig. 11a). We classified RNA samples from 20 distinct mice into these 3 distinct groups based on their ability to support long-distance CST growth. Using this approach, we could therefore eliminate differences that arose due to developmental or segmental differences and specifically identify genes that are differentially expressed between spinal levels that either do or do not support long-distance CST growth regardless of developmental time or spinal segmental level. A modified MDS plot is shown in Extended Data Fig. 11b, where the spinal samples are distinguished based on their ability to foster long-distance CST growth. We then investigated differential expression between these groups and the following comparisons were performed: 1) high vs. no long-distance growth; 2) moderate vs. no long-distance growth; and 3) high vs. moderate long-distance growth. We identified 2349 differentially expressed genes with FDR p<0.05. As expected, high vs. no long-distance growth groups showed the most difference, and fewer genes were differentially expressed between high and moderate growth groups (Extended Data Fig. 11c). The top gene ontology term that distinguishes these groups is axon guidance (Extended Data Fig. 11d). This suggests that axon guidance mechanisms represent an early control over the decline in long-distance axon growth ability in the white matter in the developing CNS.

## Discussion

Regenerative ability in the nervous system has been lost several times throughout vertebrate evolution ^29^. One approach for identifying mechanisms that result in the loss of regeneration is to examine the control over its decline through development into maturity. True and complete axon regeneration to restore function requires a transected axon to regrow after injury to its previous target(s) to reestablish the original circuitry that existed prior to injury ^13^. While this is presently an unattained goal, long-distance axon growth of an injured axon would be critical to re-establish long-range connectivity in the injured CNS ^30^. Our work enables investigation of long-distance axon growth as a specific form of axon regeneration that is distinct from other forms of axonal growth in response to injury. Hence, it is inaccurate to assume that all types of axon regeneration are governed by identical universal mechanisms. The regulation of long-distance axon growth in the white matter involves recruitment of specific additional mechanisms at the very least.

Previous investigations used neonatal lesions that disrupt axon guidance mechanisms that must be expressed in an appropriate context to direct long-distance axon growth in development. Therefore, these investigations were not able to investigate the ability of the CNS to foster long-distance axon growth. For instance, over-hemisections of the rat thoracic T8 spinal cord at P2 resulted in minimal CST growth into the lesion and not much further ^21^. Since these lesions were performed before the arrival of the CST at this spinal level, this indicates that the lack of long-distance CST growth in these instances was largely because of disruption of the normal guidance mechanisms in the spinal cord. Our results therefore strongly indicate that in contrast to these previous neonatal injury paradigms, microlesions sufficiently preserve the spinal environment such that it is not inhibitory or growth-limiting to axons. As a result, they enable more precise determination of long-distance axon growth ability through development.

Laser axotomies have been previously used to minimize perturbation of the environment to investigate axon regeneration of mammalian sensory neurons in the peripheral nervous system ^31,32^ as well as axon regeneration in other vertebrates ^33,34^ and invertebrates ^35-37^. While these studies have yielded tremendous insights into regulators of axon growth, they have been mostly limited to analyzing individual axons, which precludes their ability to investigate the fasciculated axon growth investigated in the present study. Further, for technical reasons, such approaches have not been applicable for investigating axon growth of the fasciculated axon tracts in the developing mammalian CNS. The microlesions used in this work now enables such investigation.

It was previously understood that regenerative ability declines when developmental axon growth is complete and the expression of axon growth genes that are normally highly expressed in development is downregulated in adult neurons. Misexpression of these developmental genes in adult neurons, without manipulation of the lesion environment, has been shown to increase regenerative ability of adult CSN (e.g., mTOR ^38-40^, Klf7 ^41^, Sox11 ^42^). Further, the termination of the axon growth program is followed by the formation of synapses, which also causes a decline in regenerative ability ^27^. To our surprise, we find that at P4, when at least a subset of CSN (CSN_TL_) are still in the developmental phase of axon extension to the thoraco-lumbar cord, they are still unable to mount long-distance axonal regrowth after microlesions at C2. Known developmental axon growth genes (e.g., Gap43, Cap23, Scg10, Atf3, and c-jun) are still highly expressed by CSN at this time; high expression levels by CSN are maintained at least until P14 ^43^. This indicates that high-level expression of these genes is still not sufficient to direct long-distance regrowth of these lesioned axons in the spinal white matter. In contrast, P4 microlesions at T11 showed robust long-distance axon growth to spinal levels caudal to the microlesion. At P4, only pioneer CSN_TL_ axons have arrived at T11, with ∼20% of CSN axons at thoracic T2 extending to T13 ^26^. We do not find any reduction in axon extension to the spinal levels caudal to the P4 microlesion at T11, suggesting that these pioneer axons were able to regrow after the microlesion. Furthermore, this clearly showed that the environment at the microlesion, including activated astrocytes, did not prevent late-arriving, non-lesioned axons from growing through the microlesion in their appropriate position in the spinal white matter. This is in striking contrast to previous models of neonatal spinal injuries. Astrocytic activation after more disruptive CNS lesions is known to be required for the reestablishment of tissue homeostasis ^44,45^. Given the minimal disruption caused by the microlesions, it appears consistent that we observe a very minimal astrocytic response, which is highly diminished even when compared to other models of neonatal spinal lesions ^10^.

The segmental differences in long-distance CST growth ability likely reflect a combination of mechanisms. Prior work using *in vitro* cultures of crushed neonatal spinal cords in the opossum indicates that there are segmental differences between cervical and lumbar segments that correlate with a rostral to caudal gradient of spinal cord development ^46^. These differences in regeneration were attributed to differences in levels of environmental inhibitors in the spinal cord that are known to increase with maturation ^16^. Our *in vivo* experiments, however, indicate that the distinct effects on long-distance CST growth at distinct spinal levels do not correlate with segmental differences in levels of astrocytic, microglial activation, nor with myelin levels. This is corroborated by both immunohistochemical and transcriptomic analyses. While it remains possible that there might be distinct responses by spinal cell types to microlesions at distinct spinal levels, RNA-seq analyses indicate that molecular differences between distinct spinal segments, even in non-lesioned mice, are greater than the effects of the microlesions themselves (Fig. 4). These data further corroborate that microlesions cause minimal effects on the spinal environment. Another intriguing idea is that distinct CSN subpopulations might have different responsiveness to these environmental regulators e.g., via differences in expression levels of receptors, etc. It is worth noting that even when we analyzed Crim1+ CSN_TL_, i.e., a very select CSN subpopulation, the results are identical to when we analyze the broader population. Therefore, even within very select CSN subsets, long-distance axon growth is lost at different times at different spinal levels, which argues against these spinal-level effects being CSN subpopulation specific.

One CSN-intrinsic mechanism that could account for these spinal level effects is potential differences in cytoskeletal dynamics at distinct levels along the length of a CST axon. *In vitro* evidence has identified that after an axotomy the regenerative response of a lesioned axon correlated with the length of the remaining stump – more distal cuts induced regrowth while proximal axotomies that were closer to the CSN soma transformed a dendrite into an axon. This difference was known to be controlled by microtubule stability^47,48^. Similar mechanisms could play a role in the loss of long-distance CST growth at distinct spinal levels. Using microlesions as a new approach, we can investigate whether this applies to long-distance CST growth *in vivo*.

The molecular differences between spinal segments in non-lesioned mice likely represent axon guidance mechanisms that are responsible for normally guiding CST axons to their appropriate segmental targets. These mechanisms likely also control long-distance regrowth ability. Manipulating such guidance mechanisms might help elucidate the contribution of these pathways to the initial control over long-distance regenerative ability. An implicit corollary to this idea would be that CSN-intrinsic mechanisms that normally control CSN axon growth at distinct spinal levels might also control the responses of axotomized CSN axons at distinct spinal levels. CSN projecting to distinct spinal levels express distinct genes that control their segmentally distinct axon extension ^25,26^. It is tempting to speculate that molecules that direct CSN axon extension to appropriate spinal segments would also control long-distance regrowth ability at these distinct levels. In line with this idea, we find that when CSN axons are lesioned close to the spinal level where they are normally growing (T11 at P4, T2 at P1), we find robust long-distance growth ability in the white matter. However, when the axons are lesioned at a distant site from the growing ends (C2 at P4), they show complete loss of long-distance axon regrowth. i.e., when the segmentally-appropriate axon growth mechanisms are “in sync” with where the microlesion occurs, there is robust axon regrowth. In contrast, microlesions that disrupt this developmental synchrony result in no long-distance regrowth. There is prior evidence for such “chronicity” of neuronal intrinsic mechanisms controlling regenerative ability in the developing chick CNS ^49^. In these experiments, heterochronicity of neuronal transplants shows that intrinsic developmental changes in hindbrain projection neurons are more critical than environmental changes in the spinal cord in controlling the developmental decline in the regenerative ability of these projection neurons. Therefore, control mechanisms that normally function to direct axon growth and guidance during development might also control long-distance regenerative ability. These “context-appropriate mechanisms” will include developmental stage-specific gene expression in CSN soma, appropriate and specific molecules being localized to CSN axon growth cones at the correct developmental time, and expression of correct guidance cues under spatial and temporal control in the spinal cord. It is also possible that microlesions at distinct spinal levels elicit distinct molecular responses in CSN depending on the segmental level, which could in turn underlie the spinal level-specific responses of lesioned axons. It is likely that a combination of all such mechanisms control the developmental loss of long-distance axon growth ability and future investigations will delineate their respective contributions.

Following P1 microlesions at thoracic T2, we find CSN axons maintain robust long-distance growth ability, albeit in the dorsolateral funiculus. This indicates that the signals in the spinal cord that direct long-distance CST growth are likely expressed more broadly than within a confined location in the dorsal funiculus. These results highlight that the dorsolateral funiculus is, in fact, capable of supporting long-distance growth of the majority of CST axons at least beyond thoracic T2. This further suggests largely overlapping or shared molecular mechanisms for long-distance axon growth in the dorsal as well as the dorsolateral funiculus. This finding appears consistent with the fact that CSN that normally extend axons within the dorsal funiculus versus those that extend outside it, share similar cortical locations ^50^. It remains unclear, however, whether these redirected axons after a thoracic T2 microlesion still form appropriate connectivity with their targets in the caudal thoracic and lumbar cord. Future investigations will elucidate this question.

Interestingly, even though long-distance axon growth does not occur following a P4 microlesion at cervical C2, we do observe extensive sprouting into the spinal gray matter that extends significantly further caudally into the cervical cord and largely does not extend into thoracic segments. The term “regenerative sprouting” generally describes growth arising from a lesioned axon, which does appear to apply in this case ^13,51^. It is tempting to speculate that the absence of context-appropriate growth mechanisms results in the loss of long-distance axon growth, but that the axon growth mechanisms discussed above, might be sufficient to produce such “plasticity”, i.e., sprouting.

Finally, in this work, we specifically analyzed the long-distance growth ability of the CST, but it is tempting to speculate that similar, segmentally-distinct mechanisms might also function to regulate long-distance growth of other descending, spinal-projecting pathways, e.g., rubrospinal and reticulospinal pathways. This will require experimental manipulation at earlier developmental times since their developmental growth into the spinal cord occurs *in utero* in rodents.

Together, our results provide a novel experimental approach to delineate a precise timeline of the decline in long-distance CST growth and guidance ability during development. Our results identify that this ability is regulated in a location- and time-dependent manner, and that global regulators such as astrocytic or microglial reactivity do not regulate its initial loss. The loss of long-distance CST growth can occur even when CSN are in a state of developmental axon growth, and even prior to synapse formation. Together, these results suggest that there are additional steps that control the decline of long-distance regeneration through development into maturity in a context-specific manner. Future investigations into the mechanistic underpinnings of this initial control over regenerative ability are likely to identify novel molecular substrates for corticospinal regeneration and repair following adult injury.

## Acknowledgments

We thank Sargunvir Sondhi, Jake Lustig, Aayushma Kunwar, and Alexander Lammers for their technical assistance.

This work was supported by funding from the National Institutes of Health (NIH; NINDS/ R21 NS127622, and NTRAIN/NICHD K12HD093427), a project grant from the Wings for Life Spinal Cord Foundation, a pilot grant from Craig H. Neilsen Foundation, an IDEA grant from the NY State (NYS) Spinal Cord Injury Research Board (SCIRB), and the Burke Foundation to V.S.. C.R. was partially supported by a postdoctoral fellowship from the NY State (NYS) Spinal Cord Injury Research Board (SCIRB). J.K. was partially supported by a postdoctoral fellowship from the Swiss National Science Foundation. pAAV-CAG-tdTomato (codon diversified) was a gift from Edward Boyden (Addgene viral prep# 59462-AAV1; http://n2t.net/addgene:59462;RRID:Addgene_59462)

## Author Contributions

C.R., and V.S. designed research; C.R., J.K., and P.P. performed research; C.R., J.K., F.S., and R.K. analyzed data; and C.R., and V.S. wrote and edited the manuscript.

## Methods

### Mice

All mouse studies were approved by the Weill Cornell Medical College Institutional Animal Care and Use Committee, and were performed in accordance with institutional and federal guidelines. Wild-type mice on a CD1 background were obtained from Charles River Laboratories (Wilmington, MA). The day of birth was designated as postnatal day 0 (P0). We obtained Crim1CreERT2:Emx1-IRES-Flpo:ai65 (CERai65) triple transgenic mice from Prof. Jeffrey D. Macklis where they were previously generated by crossing Crim1GCE/+, Emx1-IRES-FlpO/+, and ai65 (RCFLtdT)/+ mice ^25^. CreERT2 activity was induced as previously described whereby P3 mouse pups were injected intraperitoneally with 100 μl of tamoxifen (Sigma, T5648) solution (3.5 mg/ml) dissolved in corn oil (Sigma, C8267).

### Anterograde labeling of CST

For anterograde labeling of the CST, AAV1-CAG-tdTomato (Addgene) particles (2x 10^13 GC/ml) were injected at P0-P3 into the cortex as previously described ^25,26^. Briefly, pups were anesthetized using hypothermia for 2-3 minutes and AAV was injected unilaterally under ultrasound backscatter microscopy guidance (Vevo 2100; VisualSonics, Toronto, Canada) via a pulled glass micropipette attached to a nanojector (Nanoject II, Drummond Scientific, Broomall, PA). For mice analyzed at P35, 7x 23 nl (total volume of 161 nl) was injected, while mice analyzed at 72 h post microlesion were injected with 21 x 23 nl (total volume 483 nl). The higher titer injection for acute time points is to allow for better expression since there was less time available for AAV expression. After the injections, the pups were placed on a heating pad for recovery and returned to the dam soon after.

### Microsurgical CST lesions

Pups 0-7 days old (P0-P7) were anesthetized using hypothermia for 1.5-5 minutes. Pups older than 7 days (P8, P14, P21) were anesthetized using 2.5% isoflurane, and the fur over their backs was removed using Nair; excess Nair was removed using 70% ethanol wipes. For performing microlesions, a beveled (30°) glass micropipette (tip ∼150 μm) was used. This was attached to a high-frequency vibrating apparatus (OralB PRO 1000 electronic toothbrush, 8800 oscillations/minute, 20000 pulsation/minute). We established that the maximal displacement of the micropipette when attached to this apparatus is 400 μm. The pup was placed on its side, the dorsal side oriented towards the micropipette. The micropipette was inserted into the spinal cord at the appropriate spinal segmental level (C2, T2, or T11) through the spinal midline up to the central canal, under visual guidance provided via ultrasound-guided backscatter microscopy. Once the micropipette was in position, the vibration was turned on. During the next 10 seconds, the pipette was slowly removed from the spinal cord with brief pauses every 2 seconds to completely axotomize the CST (ultrasound video shown in Extended Data Video 1). The 10 sec period was determined as the minimum required time period to axotomize the CST (and largely similar to the 10 sec long crush lesions that were previously performed in neonatal opossums ^16^). After the microlesion, pups were placed on a heating pad for recovery and returned to the dam. We never encountered any skin lesions on the pup and there was no need for any wound closure protocol. The dams continued to take care of the pups without necessitating any additional care or intervention.

### Tissue collection and sectioning

At the experimental endpoints (72 h after microlesion for acute time points and at P35 for chronic time points), mice were transcardially perfused with cold PBS, followed by 4% paraformaldehyde (PFA) in PBS. Brains and spinal cords were carefully dissected and post-fixed in 4% PFA overnight. Prior to processing, wholemount images of the brain and spinal cord were acquired using a fluorescence stereomicroscope (Nikon SMZ18). This was to ensure that the AAV injections in all the mice were well-matched. The following blocks of tissue from each mouse were collected separately: 1) spinal cords containing the microlesion site (C1-C8 for C2 lesioned mice; C5-T4 for T2 lesioned mice; T6-L1 for T11 lesioned mice), 2) medulla for all control and lesioned groups, 3) single spinal segments for axial sections – T1 for C2 and T11-lesioned mice; T5 for T2-lesioned mice; L2 for all groups. These samples were cryoprotected separately for each mouse in 30% sucrose in 1xPBS overnight, and frozen in OCT (Tissue Tek, Sakura Finetek). All tissue blocks were cut using a cryostat (Leica CM3050S) at 50 µm (spinal cord axials) or 70 µm (spinal cord horizontals) thickness and collected as free-floating sections in 1xPBS. For horizontal sections of the spinal cord, each section was collected, stained, and mounted serially (dorsal to ventral). These were used to reconstruct the microlesion and to also analyze the placement of the microlesion site in the cord (described in greater detail below). Since serial sections were used, we were therefore able to reconstruct the microlesion volume reliably and reproducibly across the entire dorsoventral extent of the spinal cord.

### Immunohistochemistry

Non-specific binding was blocked by incubating tissue sections in 0.3% BSA (Sigma, A3059-100G) in 1x PBS with 0.3% Triton X-100. Primary antibodies were incubated in this blocking solution overnight at 4°C. The following primary antibodies and dilutions were used: mouse anti-GFAP (1:500, Sigma-Aldrich), rabbit anti-Iba1 (1:500, Fujifilm Wako), rat anti-MBP (1:500, Millipore), and rabbit anti-RFP (1:500, Rockland Immunochemicals). Next, sections were incubated with appropriate Alexa Fluor secondary antibodies for 3 h at room temperature (1:250, Invitrogen). Sections were mounted on glass slides and coverslipped using DAPI-Fluoromount-G (Southern Biotech).

### Tissue clearing and light sheet microscopy

Two whole CNS samples (brainstem+spinal cord) collected at P14 from P4 C2 and P4 T11 lesioned mice, were cleared, stained, and imaged on a light sheet microscope by LifeCanvas Technologies according to their established protocols. Briefly, the samples were collected using the same protocols as above, and after overnight fixation in 4% PFA, they were transported to the company in 1xPBS. Samples were preserved and post-fixed with SHIELD reagent, followed by clearing for 7 days in Clear+ delipidation buffer, labeled in SmartLabel with 10 μg mouse GFAP antibody. Reflective index was matched using EasyIndex solution (all reagents provided by LifeCanvas Technologies). Cleared tissues were imaged on SmartSPIM light sheet microscope at 3.6x with z-step size of 4 μm. 3D images and videos were later prepared and rendered using Imaris v.9.9.0 software (Oxford Instruments).

### Exclusion of mice from analyses

Given the small size of microlesions, we undertook stringent analyses to ensure that the site of the microlesion occurred in the correct location in the dorsal funiculus. The microlesion site was identified using GFAP immunohistochemistry to visualize the astrocytic reaction. Mice where the microlesion site was either off the midline, too ventral, or more than a segment off from the intended segmental level, were excluded. In addition, mice where AAV injections were off from the target area (caudomedial cortex) or noticeably weaker than others, were also excluded since reliable CSN axon extension analysis in these mice was not possible. We also had to exclude some mice for GFAP analyses, due to tissue folding at the lesion site. Mouse numbers for the different analyses are listed in Extended Data Table 1.

### Quantification of axon extension in the spinal cord

In mice analyzed at chronic time points (endpoint P35), axon extension in the spinal cord was quantified using 20X images of axial sections of the spinal cord taken on a Leica SP8 confocal microscope (Leica SP8) with 2 µm step z-stacks. Image analysis was done using NIH Fiji and a semi-automated custom-built macro to analyze thresholded particles in a blinded fashion. 3 sections were analyzed per mouse and the 3 brightest Z-planes from each z-stack were used for analysis. Briefly, an area containing the CST (tdTomato+) was selected as a region of interest (ROI) (pyramid in the medulla, ventral part of the dorsal funiculus, and dorsolateral funiculus in thoracic and lumbar axials), followed by manual thresholding of signal vs. background, and measurement of “Total Area of thresholded pixels” in these ROIs. Measurements at thoracic and lumbar levels were normalized to measurement at the ventral medulla and presented as a percentage for each mouse.

In mice analyzed at acute time points (72 h post-lesion), axon extension was quantified on 5x images of serial horizontal spinal cord sections containing the microlesion site, acquired on a Zeiss Axioimager M2. All sections containing labeled CST axons were used to quantify the tract in the dorsal funiculus and the dorsolateral funiculus. All analyses were performed using NIH Fiji and a semi-automated macro. In brief, dorsal and dorsolateral funiculi were outlined using DAPI images, since the spinal gray matter has higher nuclear density as compared to spinal white matter. These outlines were converted into masks and the tdTomato signal was analyzed within these masks. Every horizontal section of the cord containing the CST was then binned rostrocaudally into bins of 2000 x 200 um (W x H) extending ∼600 μm rostral, and ∼2600 μm caudal to the microlesion site. tdTomato+ pixels were manually thresholded to distinguish the signal from the background, and the number of thresholded pixels was measured in each bin for each ROI (i.e. 1 value for the dorsal funiculus and one value for each dorsolateral funiculus). The thresholded pixels in the corresponding areas and bins from each horizontal section were summed to represent the total number of CST axons present in the spinal white matter both rostral and caudal to the microlesion site for each mouse. Axon intensity bin values for individual mice were then normalized to the average CST intensity for each mouse rostral to the lesion site. Data were then presented as heatmaps of averages for each group (as shown in Fig. 4b).

### Quantification of astrocytic reactivity and its correlation to axon growth

GFAP immunohistochemistry was used to identify astrocytic activation. All analyses quantifying GFAP were performed on 5x epifluorescence images of serial horizontal sections of the spinal cord that were acquired on a Zeiss Axioimager M2. GFAP intensity at the microlesion was analyzed using NIH Fiji software. 3 sections containing the CST were chosen for each animal. The microlesion area, identifiable by increased GFAP immunoreactivity as compared to sites distant from the microlesion, was outlined and defined as the ROI. The raw intensity density (sum of the intensity values of each pixel) in each ROI was measured. We next measured GFAP intensity of an identical ROI on the same image rostral and caudal to the microlesion (baseline value), at a similar location along the midline. The intensity measured at the microlesion was then divided by the baseline value and results presented as a normalized GFAP intensity ratio for each mouse analyzed (as shown in Fig. 3b).

GFAP volume was measured using Neurolucida software (MBF Bioscience, version 2010.1.3). The lesion area (identified by increased GFAP immunoreactivity) was manually outlined in every horizontal section of the spinal cord; the experimenter was blinded to the lesion group. The dorsoventral position of each horizontal section was assigned a z-value, with adjoining sections being separated by 70 μm. Since we collected serial sections, we could then align all sections to generate a 3D model using Neurolucida (as shown in Fig. 3c-d). Neurolucida Explorer was then used to compute the full volume of the lesion area (in µm^3^).

For the correlation analyses, the volume of increased GFAP reactivity for each mouse was correlated to its respective value for axon growth (% axon area as quantified in Fig. 1e) at either T1/T5 or L2. Correlation plots are shown in Fig. 3 e,f with each dot representing an individual mouse.

Lastly, the signal distribution of GFAP (activated astrocytes) and Iba1 (reactive microglia) relative to the distance from the microlesion was quantified using a semi-automated macro written in NIH Fiji. The same sections were analyzed that were used for the GFAP intensity analysis described above. 30 ROI bins of 200 µm x 3000 µm (H x W) were drawn horizontally (rostral to caudal), covering the full width of the spinal cord using NIH Fiji. Similarly, 16 bins of 100 µm x 2000 µm (W x H) were drawn vertically (from left to right) to cover the microlesion site. 3 spinal cord sections from each mouse that contained the CST were used for this analysis. In these sections, the spinal cord outline was traced to create a mask such that we only analyzed pixels that were located within the section, and excluding the background. Then, bins of ROIs (both horizontal and vertical) were centered at the microlesion and images were manually thresholded for both GFAP and Iba1 to distinguish signal from background. For each bin, the fraction of the total area that is occupied by positive, thresholded pixels was measured to generate distribution plots as shown in Extended Data Fig. 5. Experimenter analyzing the sections was kept blinded to which lesion group the sample belonged to.

### Transcriptomics via RNA Sequencing

The dorsal spinal cord (to include both dorsal and dorsolateral funiculus) was dissected from C2, T2, and T11 spinal segments, from both non-lesioned (at P1, P4, and P7) and lesioned (72 h after P1 C2, P1 T2, P4 C2, P4 T2, and P4 T11 microlesions) pups (schematized in Fig. 5a). Each group had 4 biological replicates, with each replicate containing a dorsal spinal cord segment from one pup. RNA was extracted using Direct-zol RNA MicroPrep kit (Zymo). RNA concentration was measured on Qubit and quality confirmed using Bioanalyzer 2100 (Agilent Technologies). Subsequent steps of DNase treatment, rRNA depletion, library preparation, and sequencing were performed by Genewiz, Azenta Life Sciences. Samples were treated with TURBO DNAse to remove genomic DNA contamination, and rRNA depletion was done using QIAGEN FastSelect rRNA HMR Kit. For library preparation, NEBNext Ultra II RNA Library Preparation Kit (New England BioLabs) for Illumina was used. Next, samples were sequenced on Illumina HiSeq (2×150bp configuration, single index, per lane). Raw sequence reads were mapped to mouse mm10 genome using STAR aligner (v2.7.5c). Read counts are determined using HT-seq (v0.11.1) with mouse mm10 refSeq gene model as reference. Average input read counts were 22.4±3.3M(SD) and average percentage of uniquely aligned reads were 77.46±0.01(SD)%. Raw count matrix was filtered for low expressed genes followed by VSD normalization. Total counts of read-fragments aligned to known gene regions within the mouse ensembl gene model annotation (mm10) are used as the basis for quantification of gene expression. Fragment counts were derived using HTSeq program (v0.12.4). Ruven et al. 2023 (Preprint)

Quality control measures were performed to assess the quality of the data, including base quality, mismatch rate, and mapping rate to the whole genome. Additionally, repeats, chromosomes, key transcriptomic regions (exons, introns, UTRs, genes), insert sizes, AT/GC dropout, transcript coverage, and GC bias were assessed to identify potential issues in the library preparation or sequencing. The sequencing data have been deposited in the Gene Expression Omnibus database at NCBI (Accession GSE221353).

To identify genes that were differentially expressed between conditions, lowly expressed genes were removed and genes with counts per million (CPM) greater than 1 in at least 4 samples were kept for downstream analysis. We used DESeq2 package, R version 4.1.1. for TMM normalization. TMM-normalized count matrix was then used for principal component analysis. To remove developmental and section effects, Remove Unwanted Variation (RUV) ^52^ with k=4 was used. Differentially expressed genes were then determined using the Bioconductor package EdgeR, (ver 3.14.0) at False Discovery Rate (FDR) of 5%. Furthermore, GO term enrichment analysis was performed using hypergeometric overlap test.

### Statistical analysis

For the comparison of multiple groups, we used one-way ANOVA followed by Tukey’s posthoc test where microlesion groups were compared to the control, non-lesioned mice. A linear regression model was employed to assess the potential correlation between lesion volume and axon area, with the resultant R-squared value serving as an indicator of the observed relationship (R^2^ value of 0 indicates no correlation). All statistical tests were performed, and graphic presentations obtained using Graphpad Prism 9.2. A p-value of <0.05 was considered statistically significant. No statistical methods were used to pre-determine sample sizes.

---

**Extended Data Fig. 1.**
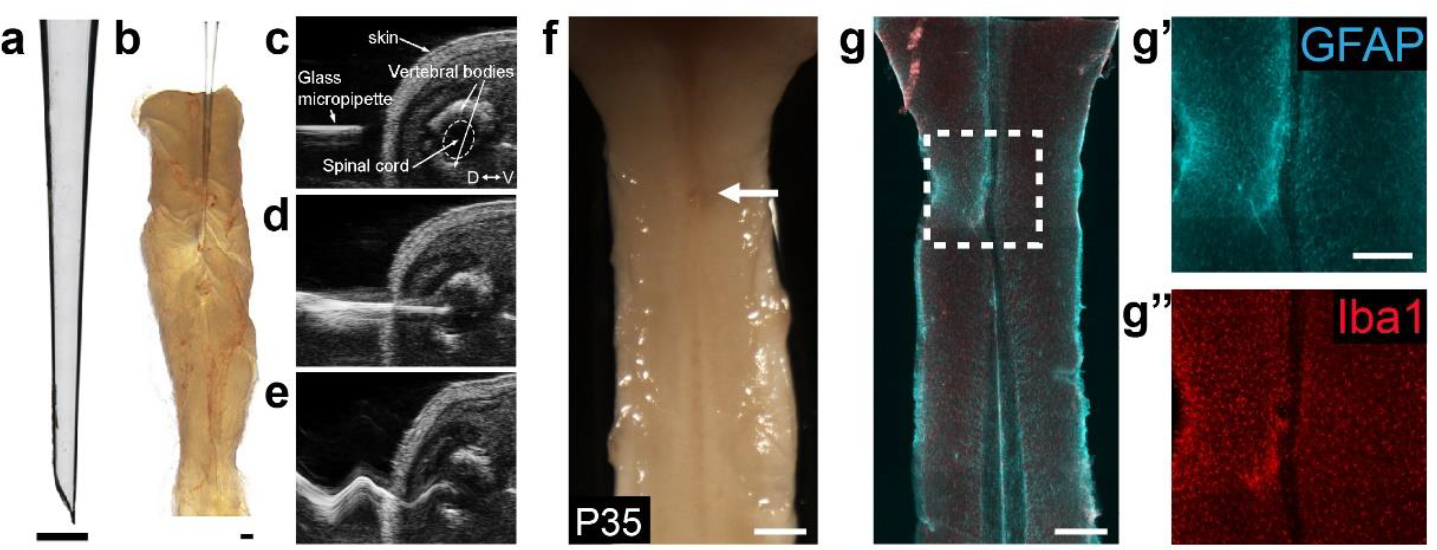
A novel approach for microsurgical CST lesions. **a**. Beveled glass micropipette used for microlesion. **b**. The same micropipette is seen inserted into a neonatal spinal cord *ex vivo*. **c-e**. Ultrasound images of the microlesion protocol show the micropipette prior to insertion into the spinal cord (**c**), after insertion into the spinal cord (**d**), and after the vibrating apparatus has been activated to produce the microlesion (**e**). **f**. Cervical cord from a P35 mouse after microlesion at P4 (arrow indicates microlesion site). **g-g”**. Horizontal section of the same cord stained for GFAP (astrocytes) (**g’**) and Iba1 (microglia) (**g”**) shows minimal astrocytic and microglial activation at the microlesion. Scale bars: **a, b, g’**: 250 μm; **f, g**: 500 μm.

**Extended Data Fig. 2.**
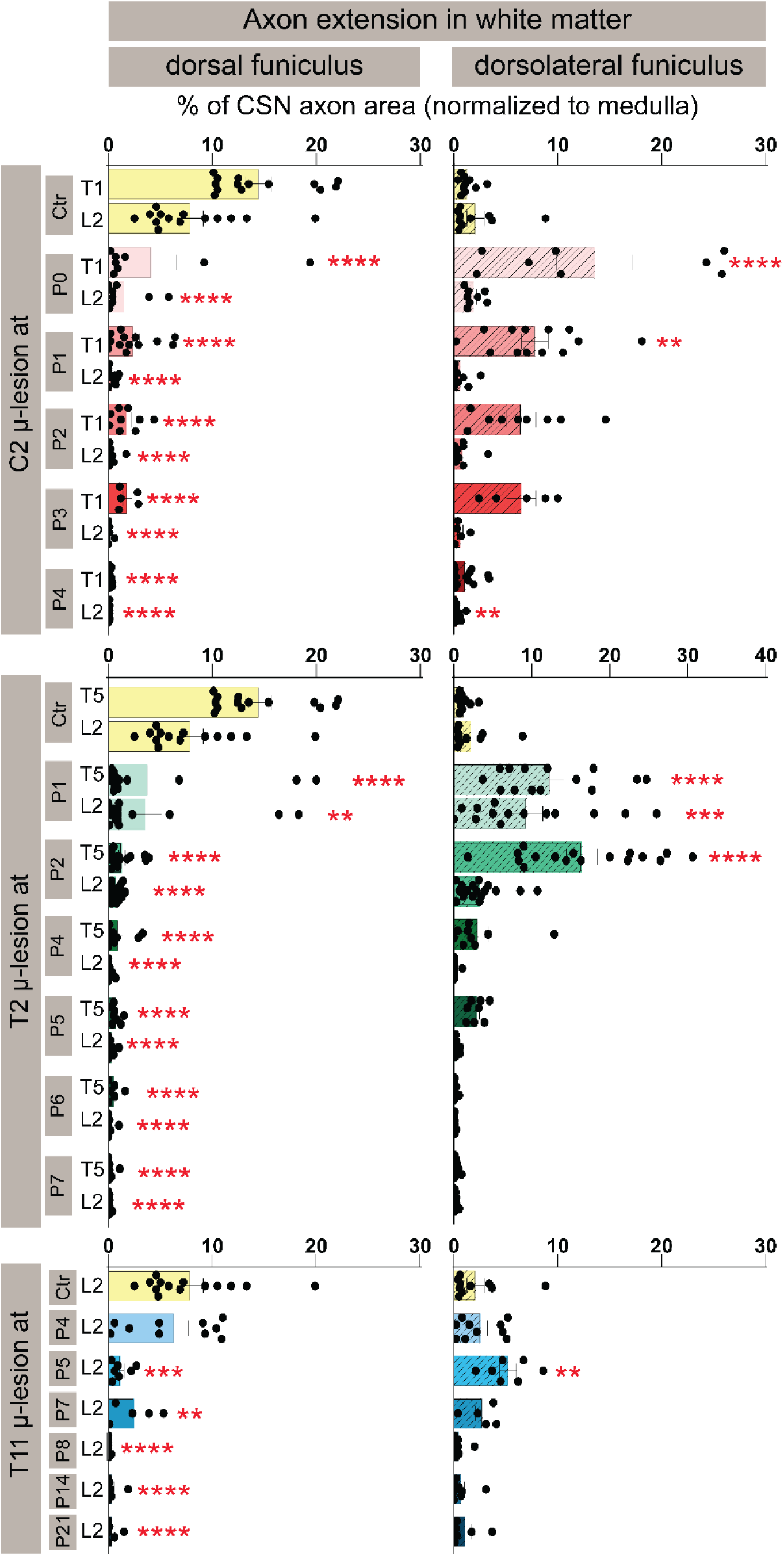
CST axons get diverted from dorsal funiculus to dorsolateral funiculus in distinct microlesioned groups. Quantification of CST extension in the spinal white matter after distinct microlesions. Thresholded area of CST axons at thoracic T1/T5 and lumbar L2 (normalized to the total area in ventral medulla) is shown separately for the dorsal and dorsolateral funiculus. * p<0.05, ** p<0.01, *** p<0.001, **** p<0.0001 injured DF/dLF vs. control DF/dLF. Data are mean + s.e.m.. Each dot represents a separate mouse.

**Extended Data Fig. 3.**
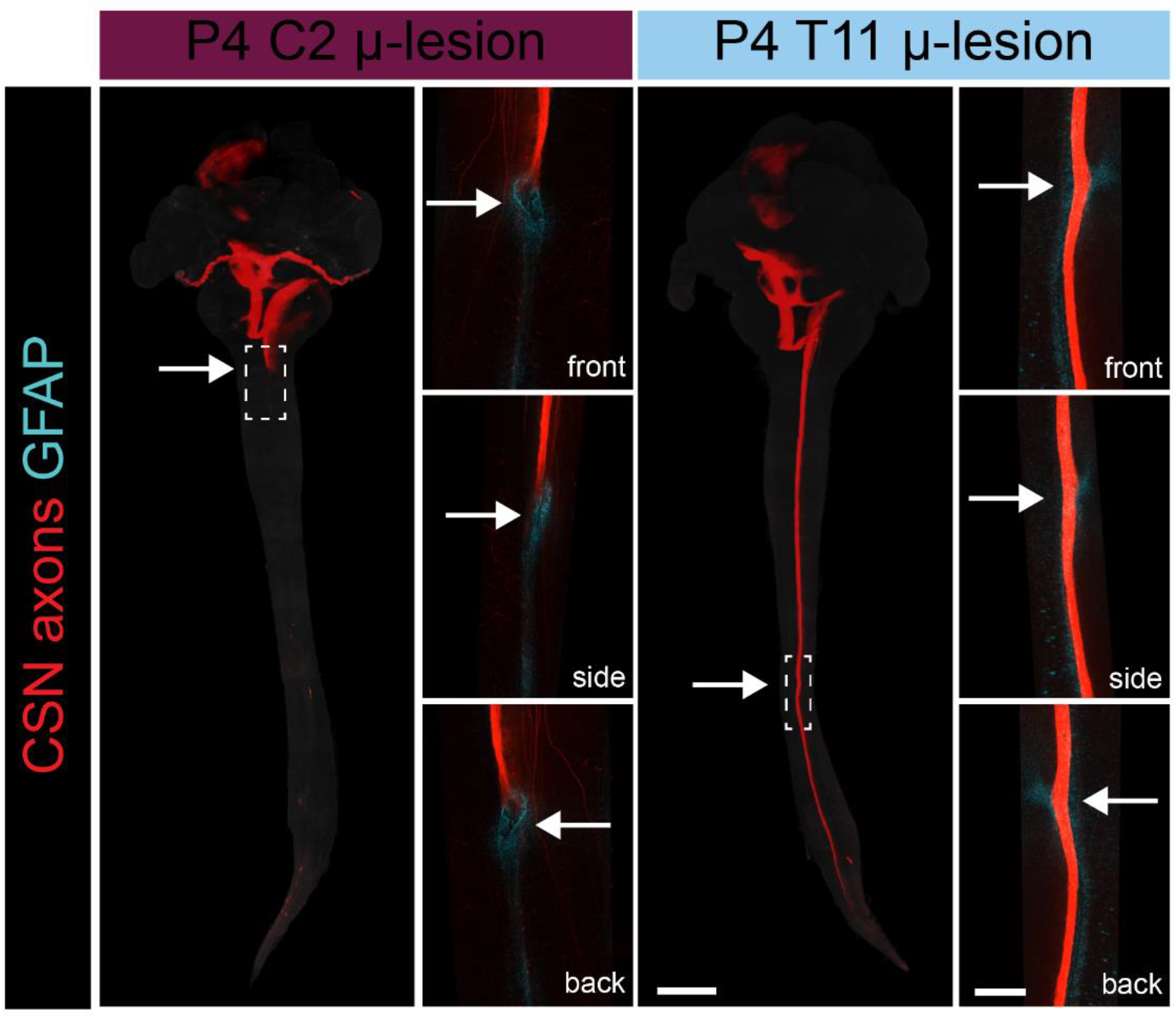
Optically cleared tissue samples highlight differential loss of long-distance growth ability at distinct spinal segments. The brainstem and spinal cords from mice that underwent P4 microlesions were optically cleared at P15 and imaged using light-sheet fluorescence microscopy. CSN axons (in red) do not extend past a P4 C2 microlesion (white arrow) but do extend past a P4 T11 microlesion. Closeups are showing stopped or growing CSN axons in relation to the lesion site (defined by the area of reactive astrocytes, GFAP+ in cyan). Scale bars: 2,000 μm full spinal cord, 500 μm closeups.

**Extended Data Fig. 4.**
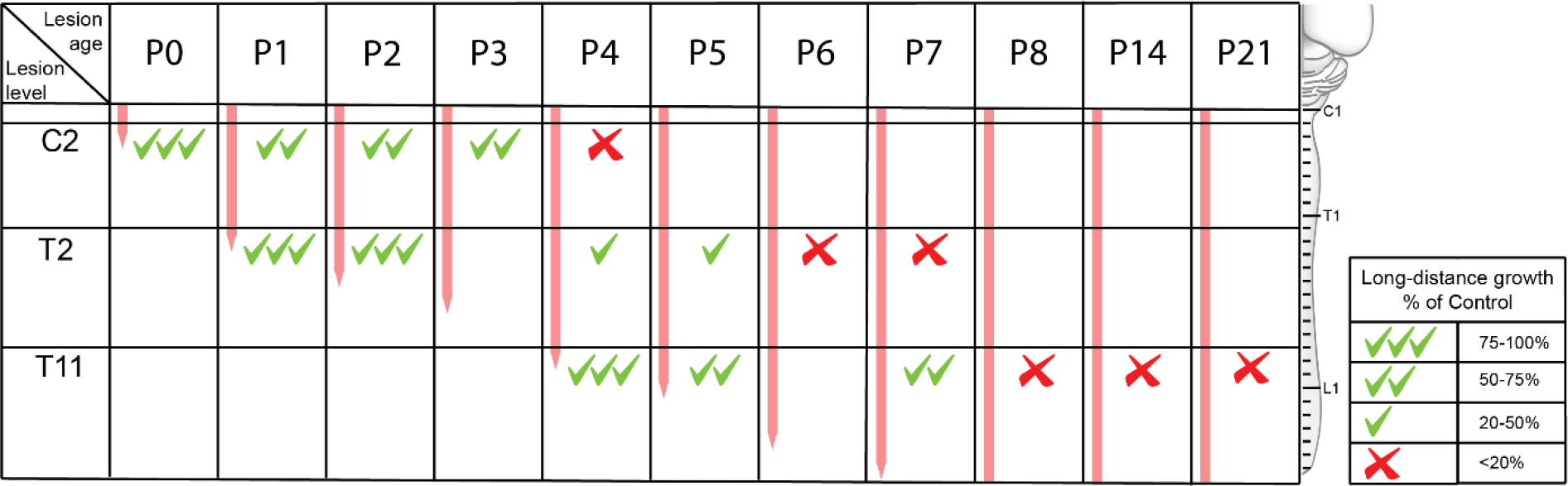
Spatio-temporal trajectory of loss of long-distance CST growth ability during development. Summary schematic shows the long-distance CST growth ability at cervical C2, thoracic T2, and thoracic T11 at distinct developmental times. The segmental level where the growing CST axons are present at the time of the microlesion is indicated by the level of the red line (adapted using data from Bareyre et al 2005). Green tick marks indicate the presence of long-distance growth ability; the number of tick marks represents the robustness of the ability at that level (see graded scale on the right). Red “x” denotes the loss of long-distance CST growth ability. Long-distance growth ability is calculated as a percentage of control axonal area at T1 for C2 microlesions, at T5 for T2 microlesions and at L2 for T11 microlesions. At cervical C2, the leading ends of the CST have reached this segment by P0. At this time, long-distance growth ability is intact. The ability declines from P1 to P3 at which times there is modest long-distance growth, and the ability is abolished by P4. At thoracic T2, long-distance growth ability remains high at P1 and P2, declines to moderate levels until P5, and the ability is abolished by P6. At thoracic T11, the ability remains fully intact at P4 and is lost by P8.

**Extended Data Fig. 5.**
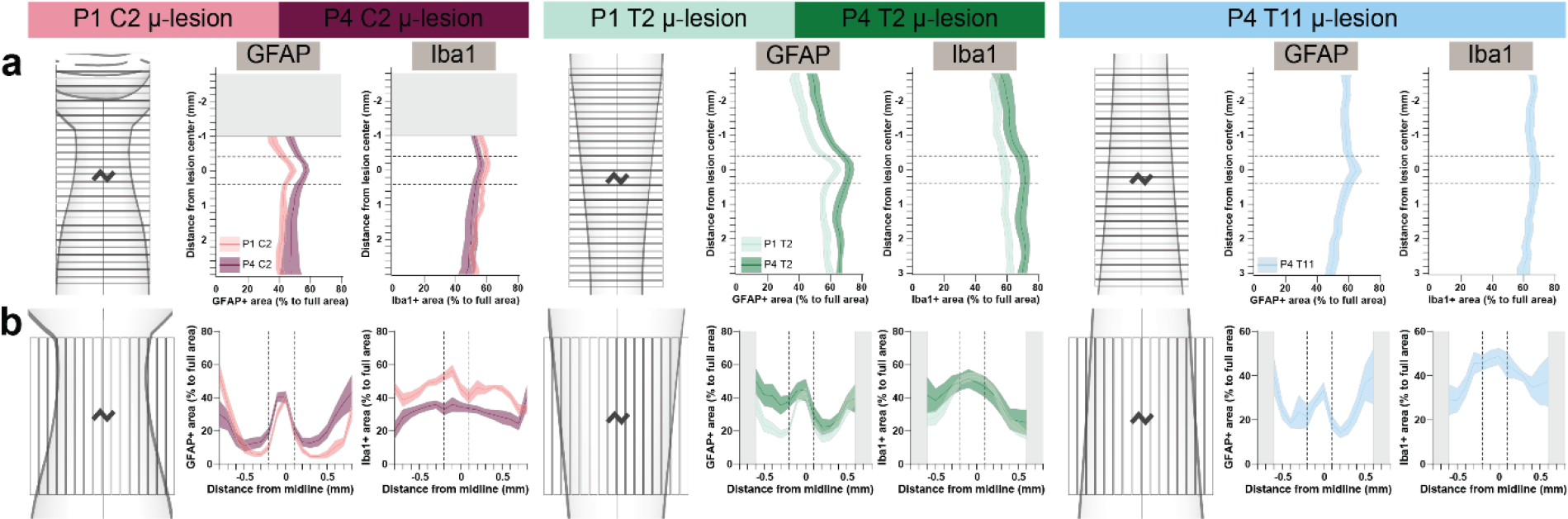
Segmentally distinct loss of long-distance CSN axon extension is not due to segmental differences in astrocytic or microglial activation. a-b. Quantification of GFAP and Iba1 distribution relative to the distance from the microlesion in P35 spinal cords that underwent microlesions at the different levels and developmental times indicated. Signal distribution was quantified in either 200 μm horizontal bins (rostral to caudal) (**a**) and 100 μm vertical bins (left to right) (**b**) as shown in the schematics. The kinked line in the schematics represents the microlesion. Data are mean ± s.e.m. The extent of the microlesion is marked with horizontal (**a**) and vertical (**b**) dashed lines on each graph. There is no difference in the overall spatial distribution of GFAP or Iba1 activation between the multiple, distinct groups.

**Extended Data Fig. 6.**
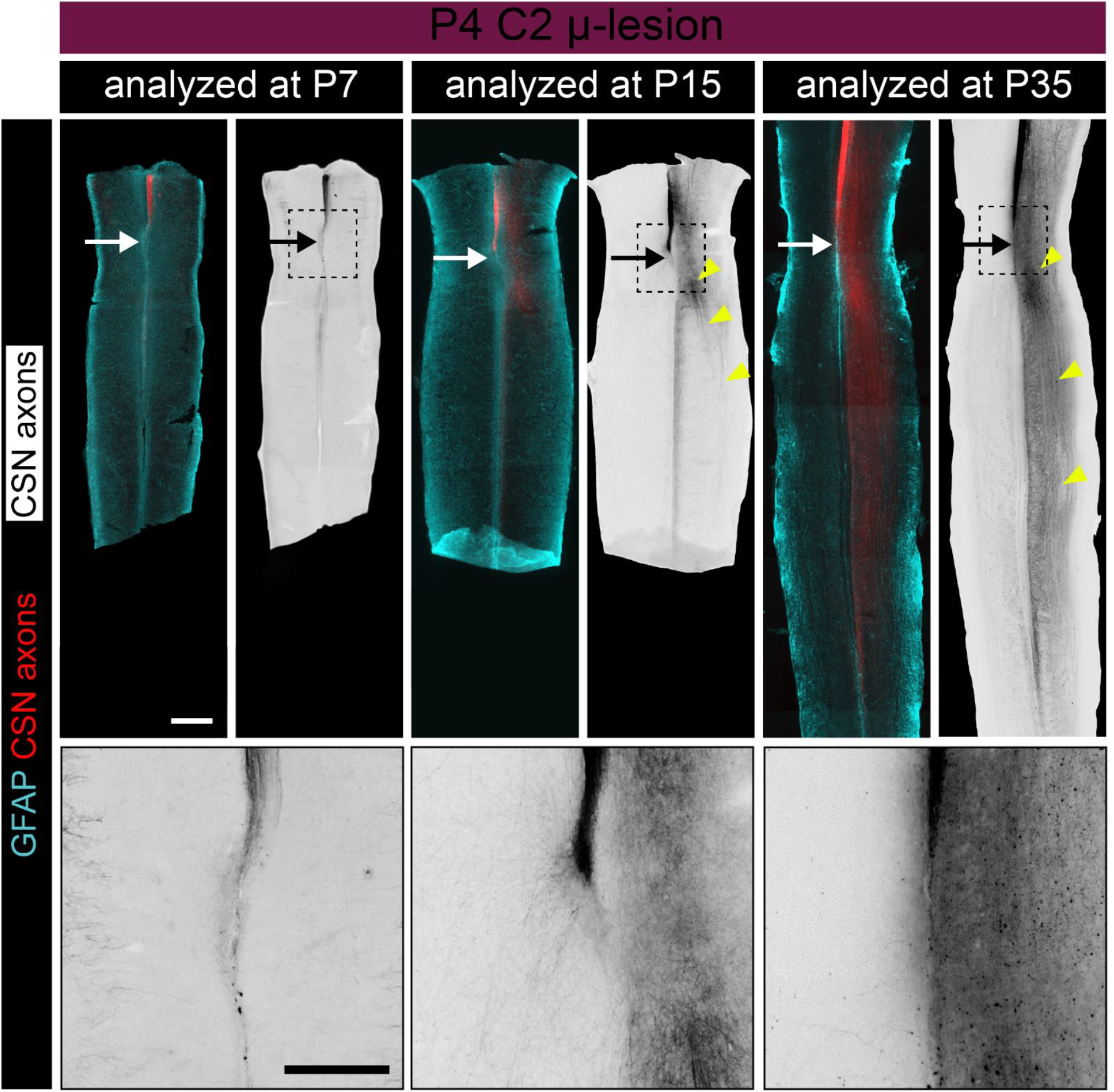
Axons retain the ability for sprouting in the gray matter even when their ability for long-distance growth in the white matter is lost. Horizontal spinal cord sections from mice that underwent a P4 C2 microlesion. Spinal cords were imaged at P7 (72 h after microlesion), P15 (11 days after microlesion), and P35 (31 days after microlesion). GFAP+ astrocytes (cyan) delineate microlesion (white arrow); CST axons are in red. Monochrome images in the adjacent panels also show CSN axons. The lack of axon growth is evident at P7, at which time the CST is seen halted at the site of the microlesion. At this time, there is no growth in the gray matter. By P15, while CST axons do not extend in the spinal white matter, they have begun to sprout into the gray matter (yellow arrowheads). This sprouting into the gray matter, which is reminiscent of regenerative sprouting, increases by P35. Lower panel shows the closeups of the injury site. Scale bars: 500 μm, closeups: 250 μm.

**Extended Data Fig. 7.**
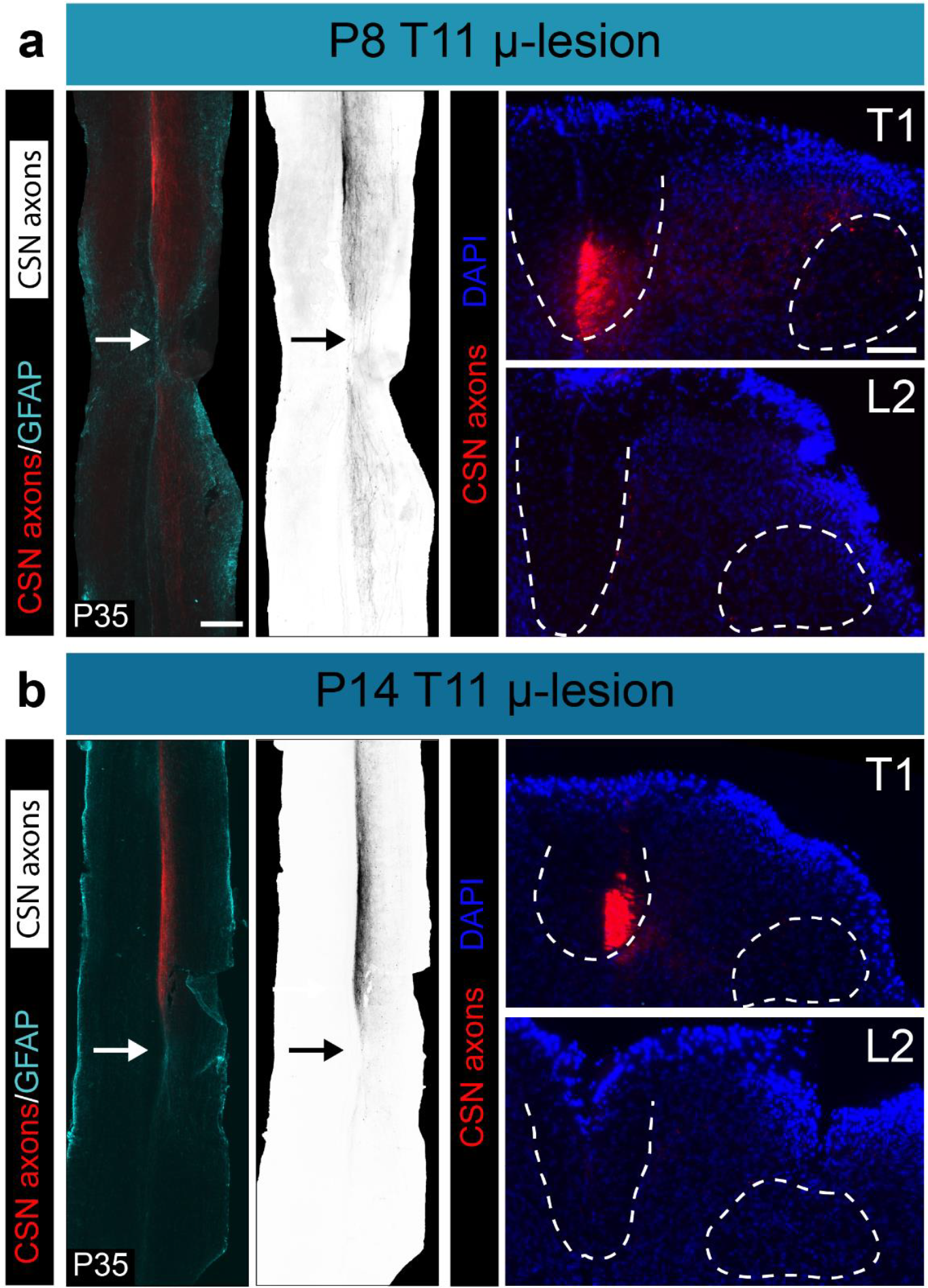
Axonal ability for growth in the gray matter declines days after the ability for long-distance growth is lost. **a**. Left: Horizontal section of a P35 thoracic spinal cord from a mouse that underwent a P8 T11 microlesion. CSN axons are in red; GFAP in cyan. Monochrome image shows only CST axons. Arrows indicate the microlesion. Right: Axial sections from the same mouse at thoracic T1 and lumbar L2. There is no long-distance CST growth in DF or dLF (demarcated by dotted outlines); however, numerous axons enter the gray matter extending caudal to the level of the microlesion. **b**. Left: Horizontal section of a P35 thoracic spinal cord from a mouse that underwent a P14 T11 microlesion. CSN axons are in red; GFAP in cyan. Monochrome image shows only CST axons. Right: Axial sections from the same mouse at thoracic T1 and lumbar L2. There is no long-distance CST growth in DF or dLF (demarcated by dotted outlines). Unlike the P8 microlesioned mouse, very few axons sprout into the grey matter after a P14 microlesion. Scale bars: **a, b**: 500 μm in horizontal sections, 100 μm in axial sections.

**Extended Data Fig. 8.**
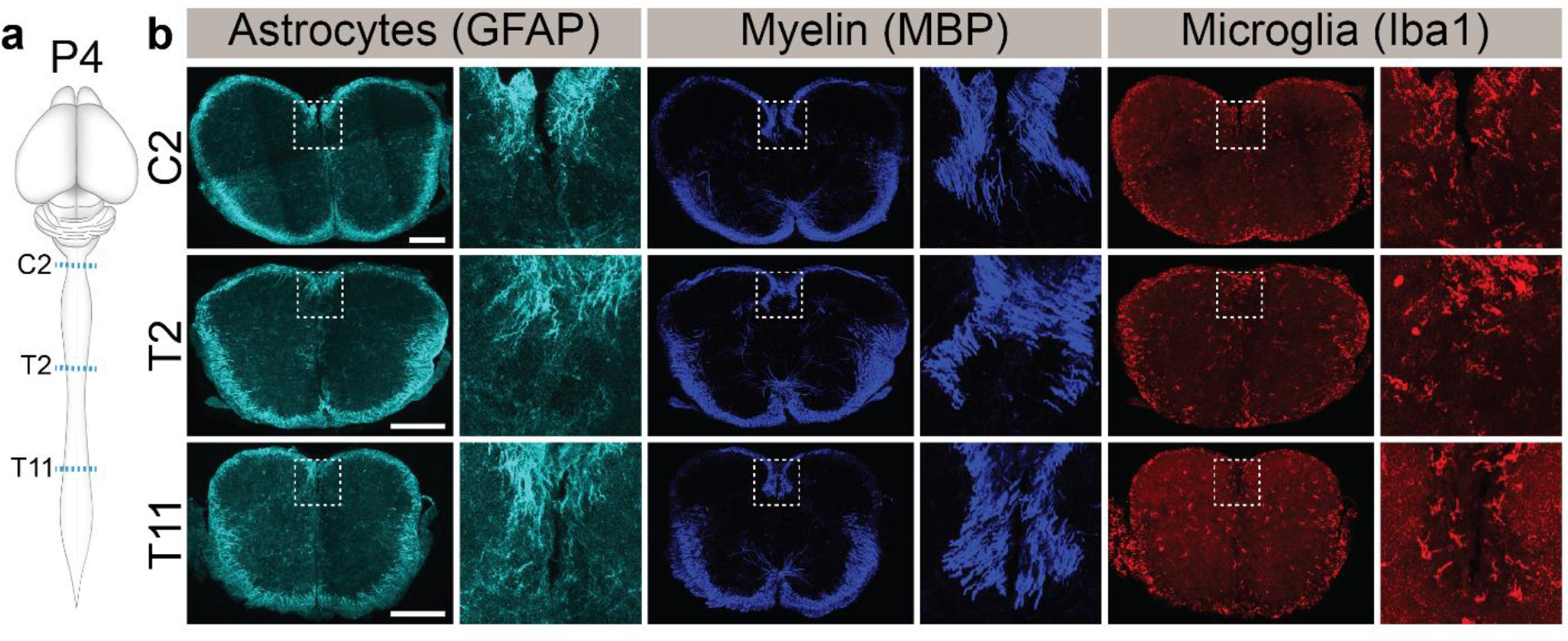
No discernible differences in astrocytes, oligodendrocytes, and microglia at distinct spinal segments in P4 non-lesioned mice. **a**. Schematic shows the spinal levels (C2, T2 and T11) of axial sections taken from P4 non-lesioned mice. **b**. Axial sections immunolabeled for astrocytes (GFAP, cyan), myelin (MBP, blue) and microglia (Iba1, red); insets show higher magnification images of the dorsal funiculus (boxed area demarcated by dotted outlines). There are no overt differences between the three segments. Scalebar: 200 μm.

**Extended Data Fig. 9.**
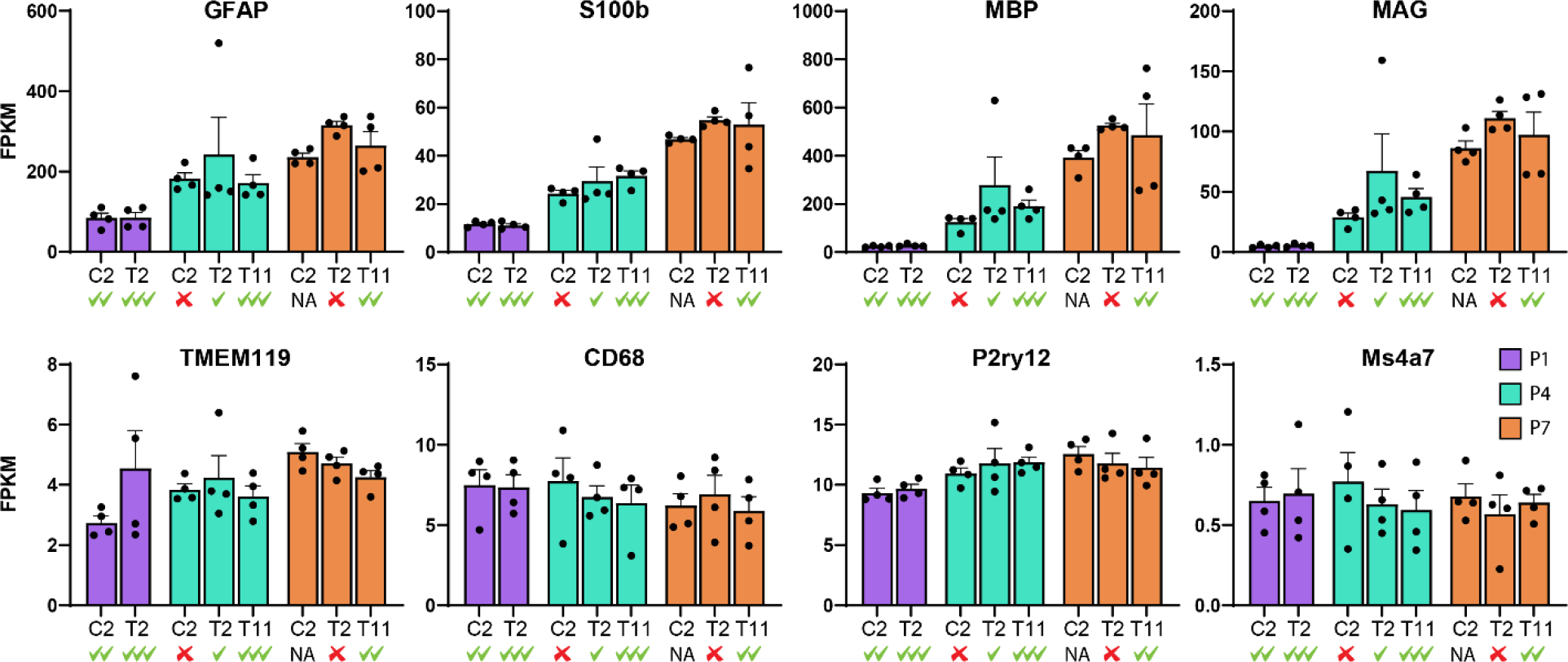
Temporal changes in astrocytes, myelination and microglia during development do not correlate to the axon growth ability. RNASeq of the dorsal cord from non-lesioned animals at various ages and spinal levels. There is increase in astrocytes and myelination from P1 to P4 to P7 that does not correlate with the ability of that spinal segment to foster long-distance growth when a microlesion occurs at these timepoints at these levels (√s indicate intact long-distance growth ability while x indicates no ability; inferred from primary quantification in Figure 1 and as schematized in Extended Data Fig. 3). Microglial gene expression does not show significant temporal or spatial differences in the non-lesioned cord. Gene expression data is presented as FPKM and shown as mean + s.e.m.. Each dot represents a single mouse; n=4 mice for each bar graph.

**Extended Data Fig. 10.**
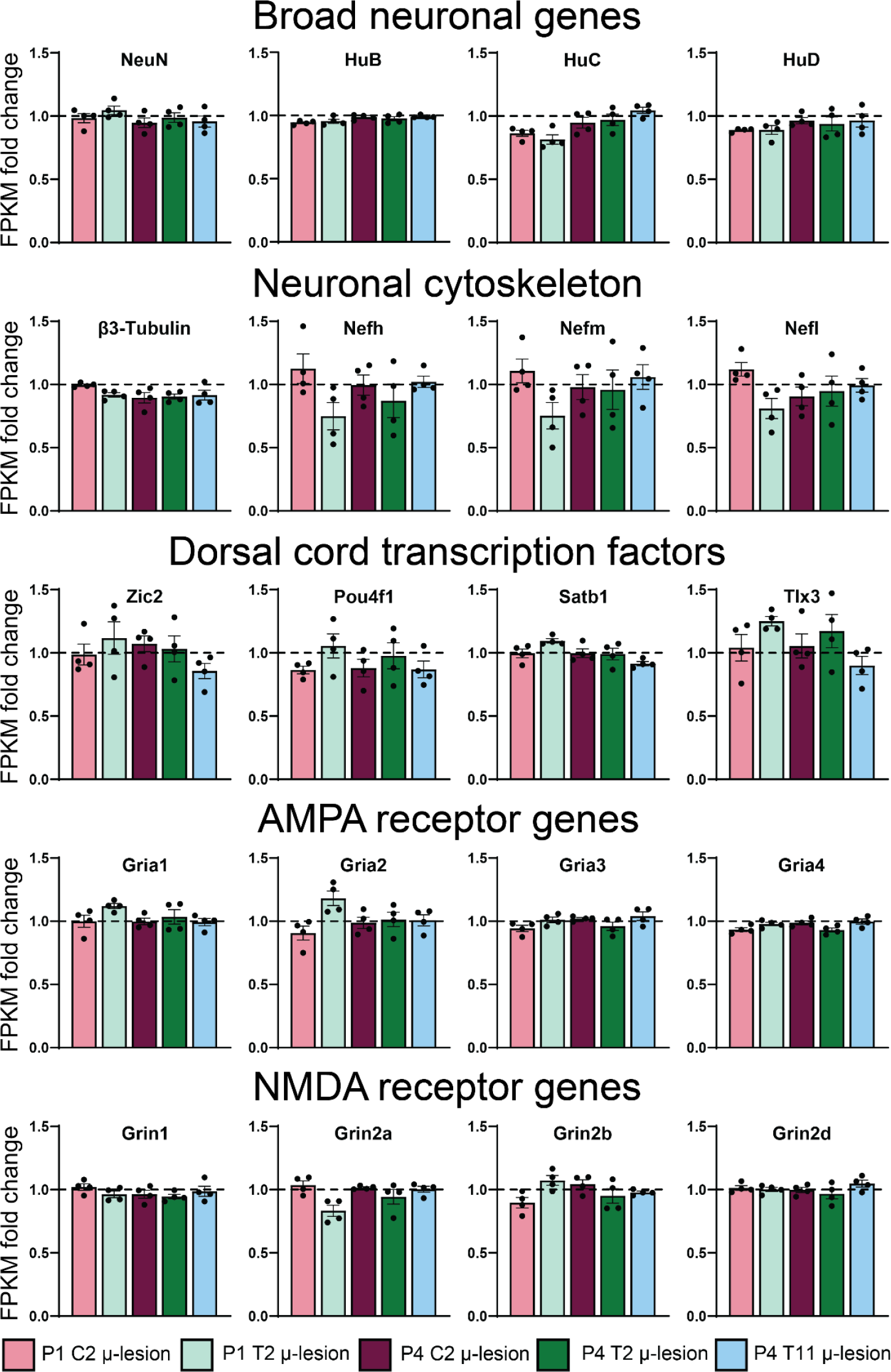
Microlesions do not overtly disrupt spinal cord. RNASeq of the dorsal cord from mice microlesioned at various ages and spinal levels. There is no difference in expression levels between any of the microlesioned groups. Expression levels in microlesioned mice are plotted as fold change over gene expression in control non-lesioned mice at the corresponding developmental time and spinal level. All expression data is presented as FPKM and shown as mean ± s.e.m..

**Extended Data Fig. 11.**
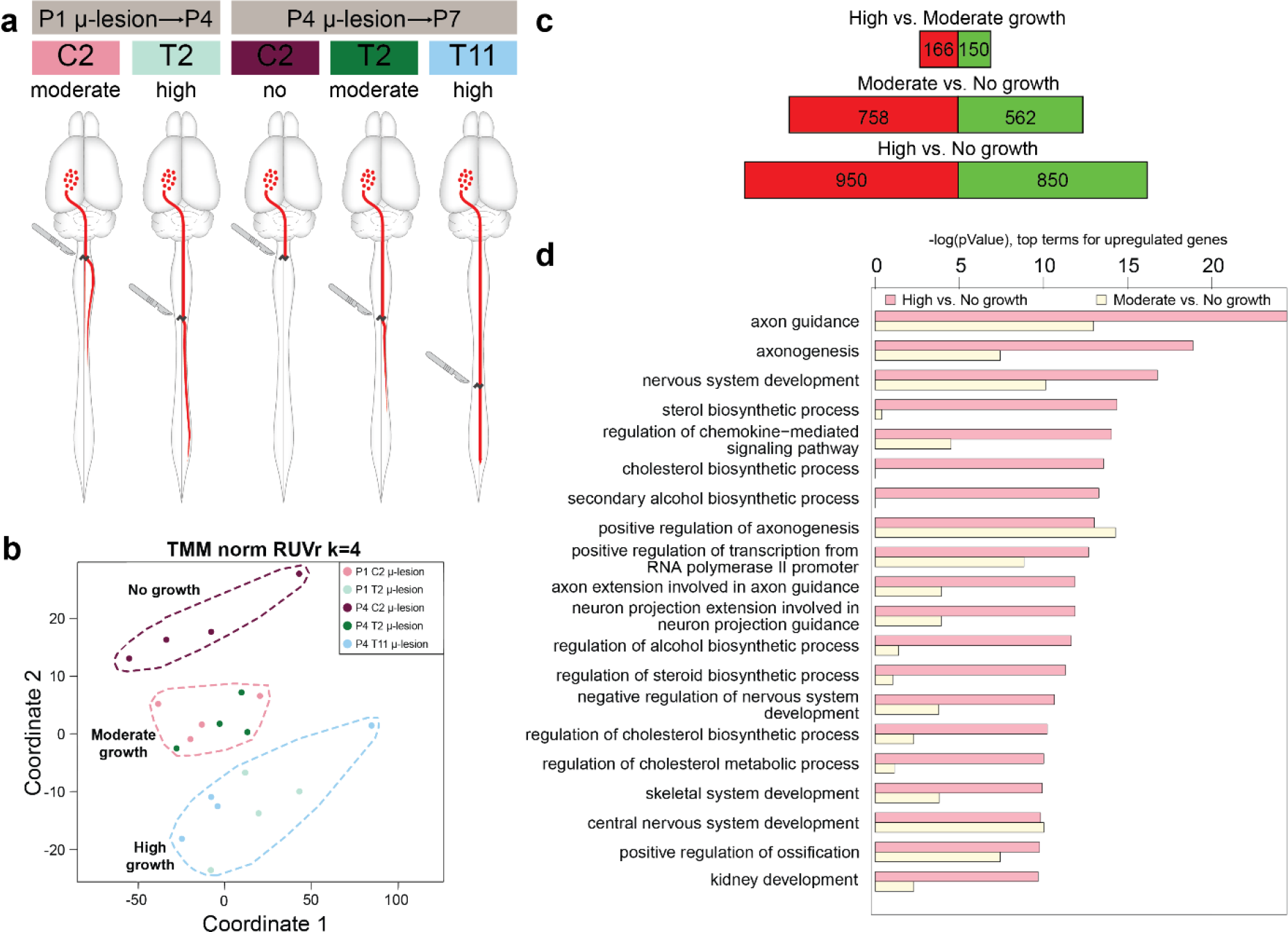
Transcriptional differences that delineate long-distance growth ability. **a**. Schematic summarizing loss of long-distance CST regenerative ability in development. At P1 (control), the growing ends of the CST are at thoracic T3. P1 microlesions at thoracic T2 result in robust long-distance axon growth (diverted from dorsal to dorsolateral funiculus); moderate long-distance growth ability is present at cervical C2. At P4 (control), the growing ends of the CST are at thoracic T13, and P4 microlesions at 3 distinct spinal segments produce distinct effects on long-distance CST growth – the ability is almost completely lost at cervical C2, moderately intact at thoracic T2, and fully intact at thoracic T11. **b**. Updated MDS plot of micro-lesioned spinal samples that distinguishes them on the basis of their ability to support long-distance CST growth into high, moderate, and no long-distance growth groups (details in text). **c**. Number of differently expressed genes between high, moderate and no long-distance growth groups. Red bars indicate the number of downregulated and green bars the number of upregulated genes. **d**. Gene ontology term enrichment analysis shows the top 20 terms that were found in the comparison of High vs. No growth groups (pink bars). In addition, values for Moderate vs. No growth are shown (light yellow bars). The GO term “axon guidance” is the term with the highest significance (-log(pValue) on the horizontal axis).

**Extended Data Video 1.**
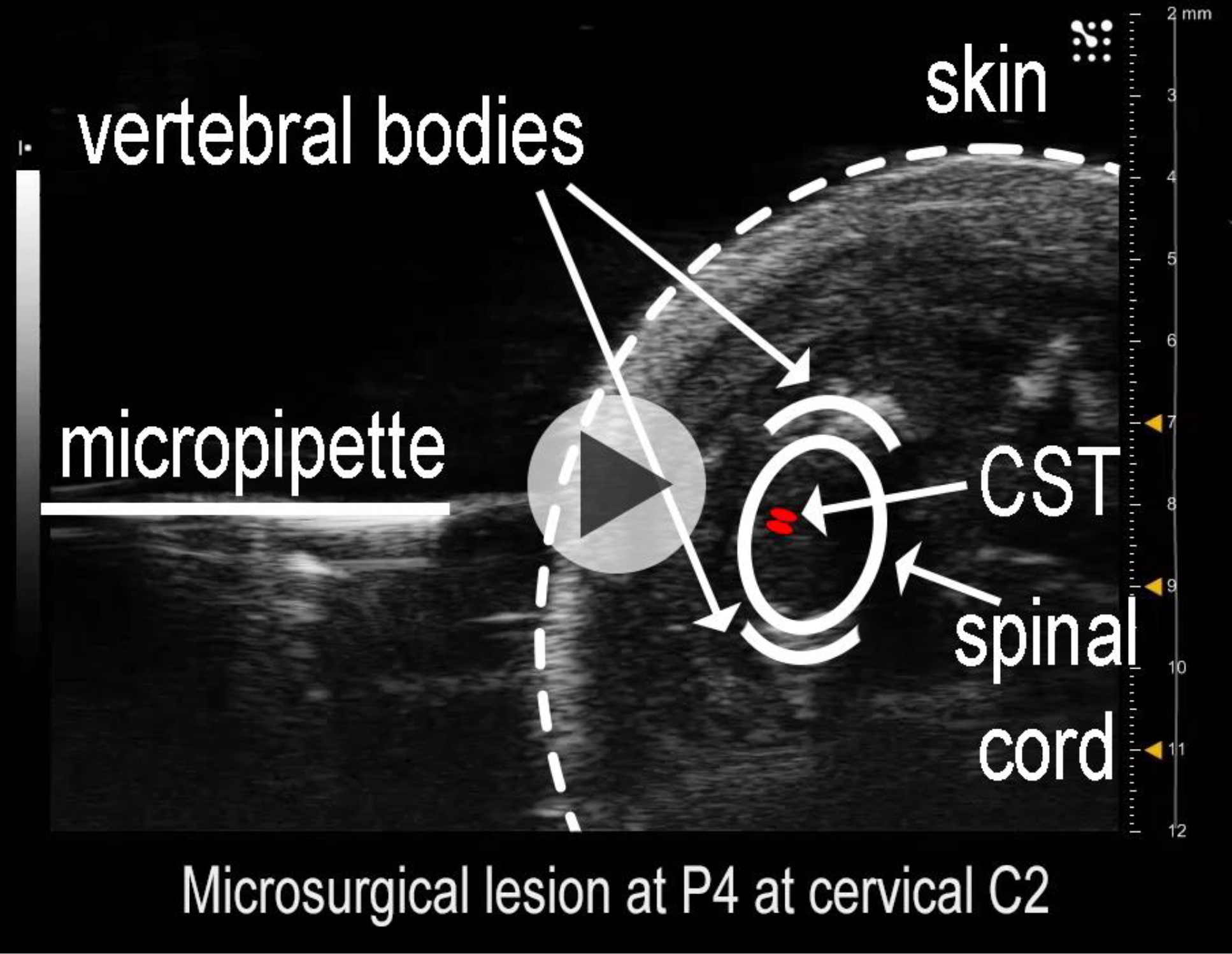
A novel approach for microsurgical CST lesions (“microlesions”). A video taken under ultrasound-guided backscatter microscopy showing the microlesion procedure. An axial view of a P4 mouse pup on its side, dorsal side towards the micropipette (on the left). Spinal cord, vertebral bodies and CST are highlighted in the initial frames of the video for reference. In the video, the glass micropipette is first seen prior to insertion into the spinal cord. It is then inserted up to the central canal, and the high-frequency vibrating apparatus is activated to axotomize the CST. After the activation of the vibration, the micropipette is withdrawn from the cord over a duration of 10 seconds (with brief 2 s pauses during the withdrawal).

**Extended Data Video 2.**
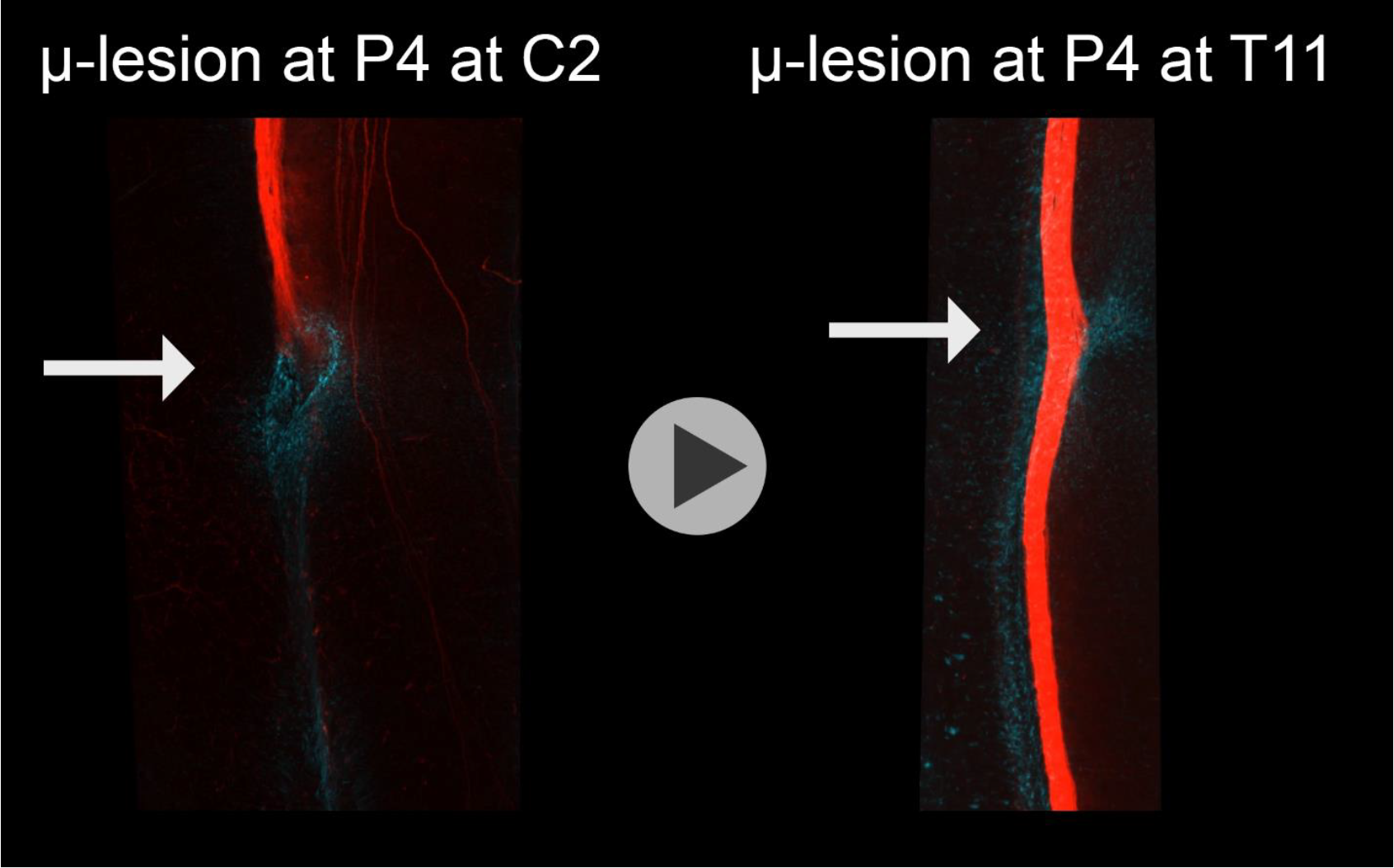
Distinct lesion responses at P4 C2 and P4 T11 microlesions. Videos showing a 3D view from an optically cleared spinal cord after either a P4 C2 or P4 T11 microlesion. The video shows a closeup view of the microlesion site (arrow; reactive astrocytes in cyan and CSN axons in red). Note that CSN axons do not extend past the P4 C2 microlesion but extend past the P4 T11 microlesion.

**Extended Data Table 1.** Animal numbers used for various analyses.

| Analysis |  | Axon extension | GFAP intensity | GFAP volume | GFAP distribution |
| --- | --- | --- | --- | --- | --- |
| Figure Group |  | Fig. 1e<br>Ext. Data Fig. 2 | Fig. 3b | Fig. 3d-f | Ext. Data Fig . 8 |
| Non-lesioned |  | 14 | NA | NA | NA |
| Age and level at which microlesion was performed | P0 C2 | 8 | 8 | 5 |  |
|  | P1 C2 | 13 | 13 | 13 | 10 |
|  | P2 C2 | 9 | 9 | 9 |  |
|  | P3 C2 | 5 | 5 | 5 |  |
|  | P4 C2 | 14 | 13 | 14 | 13 |
|  | P1 T2 | 14 | 14 | 14 | 14 |
|  | P2 T2 | 17 | 16 | 16 |  |
|  | P4 T2 | 9 | 9 | 9 | 9 |
|  | P5 T2 | 8 | 8 | 8 |  |
|  | P6 T2 | 6 | 5 | 4 |  |
|  | P7 T2 | 9 | 6 | 5 |  |
|  | P4 T11 | 9 | 9 | 8 | 9 |
|  | P5 T11 | 7 | 7 | 7 |  |
|  | P7 T11 | 5 | 5 | 5 |  |
|  | P8 T11 | 8 | 0 | 0 |  |
|  | P14 T11 | 8 | 8 | 8 |  |
|  | P21 T11 | 6 | 6 | 6 |  |
| Total |  | 169 | 141 | 136 | 55 |

